# Cell DiffErential Expression by Pooling (CellDEEP) highlights issues in differential gene expression in scRNA-seq

**DOI:** 10.64898/2026.03.09.710522

**Authors:** Yiyi Cheng, Toby Kettlewell, Ross F Laidlaw, Olympia M Hardy, Andrew McCluskey, Thomas D Otto, Domenico Somma

## Abstract

Accurate identification of differentially expressed genes (DEGs) in single-cell RNA sequencing (scRNA-seq) data remains challenging. Single-cell–specific statistical models often report large numbers of candidate genes but can exhibit inflated false positive rates, whereas pseudobulk approaches improve false discovery control at the cost of reduced sensitivity. To overcome the noise and bias that other tools have, and allow the user to have more control of the DEG process, we present CellDEEP, which uses a cell aggregation (metacell) approach. This tool provides a framework for flexible selection of pooling strategies and parameterisation for differential expression analysis (DE).

Benchmarking on simulated and real datasets, including COVID-19 and rheumatoid arthritis, shows that CellDEEP often outperforms other methods, consistently reduces false positives compared to single-cell methods and recovers more true positives than pseudobulk methods. Our work shifts the focus from selecting a single “best” method to an approach that reduces cell-level noise while preserving biological signal, together with transparent validation framework, advancing more reliable differential-expression analysis in single-cell transcriptomics.

**Graphical abstract:** 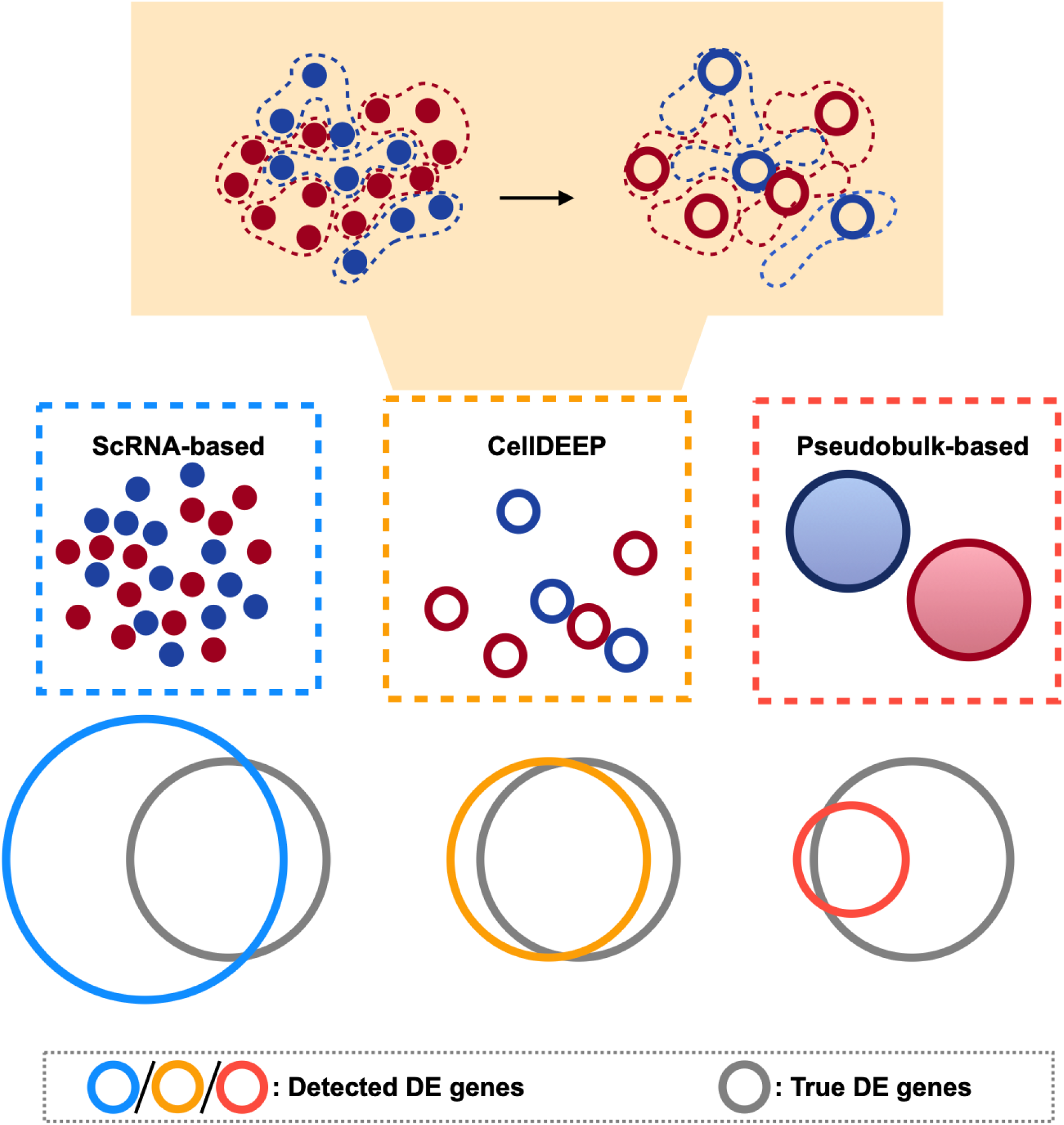

## Introduction

RNA sequencing (RNA-seq) has transformed transcriptomic studies, enabling the exploration of gene expression across tissues and conditions. Bulk RNA-seq, which measures average expression across pooled cells, has been instrumental in disease classification and response prediction, but it obscures cell-specific dynamics by averaging expression across diverse cell types. In 2009, single-cell RNA sequencing (scRNA-seq) emerged^1^ and gained traction as the cost of high-throughput platforms declined. It enables the detection of rare cell types, lineage tracing, and gene regulation^2^. However, this power comes at a cost: scRNA-seq data are inherently sparse, with high dropout rates and zero-inflation, in which genes may be undetected despite being expressed, posing significant challenges for differential expression (DE) analysis^3,4^.

Established DE methods for bulk RNA-seq, such as DESeq2^5^, edgeR^6^, and limma^7^, model expression using statistical distributions (e.g., negative binomial) and can account for complex experimental designs through generalised linear models (GLMs). For scRNA-seq, these methods are often applied using a pseudobulk approach, where expression counts are aggregated across cells from the same condition and sample^8^. While this approach improves statistical power and controls false positives, it sacrifices cell-level resolution.

Alternative scRNA-seq–specific methods such as MAST^9^ model dropout explicitly and retain single-cell resolution, offering improved sensitivity in detecting subtle gene-expression changes. Yet these methods can inflate false positive rates, particularly when the assumptions about data distribution do not hold^10^.

The performance of DE methods varies widely across benchmarking studies. Pseudobulk methods are generally preferred for their robustness^4,11^, while scRNA-based methods offer higher sensitivity in contexts where capturing subtle differences is crucial^12^. It has also been proposed that the DE strategy should be tailored to the study design: for case-control studies, pseudobulk methods are preferable, whereas for intra-condition comparisons (e.g., developmental trajectories), scRNA-based methods may be more suitable^13^.

Recently, linear mixed models for scRNAseq data have been developed. While promising, it is unclear how widely they are adopted, and they face practical challenges: benchmarks indicate they do not consistently outperform the simpler, well-established pseudobulk approaches, and they can be computationally intensive^11,14^.

Motivated by these limitations, we developed CellDEEP (Cell DiffErential Expression by Pooling), a hybrid DE analysis framework that aims to combine the strengths of both approaches. By selectively pooling cells prior to DE testing, CellDEEP reduces noise and zero inflation while retaining resolution and biological signal. In this study, we benchmark CellDEEP against scRNA-based and pseudobulk-based DE methods, using both simulated and real-world datasets, evaluate the effects of pooling strategies and data quality, and provide practical guidance for differential expression analysis in scRNA-seq studies.

## Methods

### CellDEEP architecture

CellDEEP is designed to merge single cells into metacells before DE analysis to reduce the likelihood of false positives in the standard scRNAseq DE analysis pipeline. The complete pipeline consists of three main steps: *dataset preprocessing, metacell creation, and DE analysis*.

**Dataset preprocessing** extracts three key identifiers required for downstream analysis: group ID, sample ID, and cluster ID.

The **metacell creation** workflow consists of three main steps: (i) defining subset datasets, (ii) selecting cells for pooling, and (iii) aggregating gene read counts. Cells are first separated into multiple subset datasets based on shared cluster, group, and replicate labels. Each cluster *k*, group *g*, and replicate *r*, we define

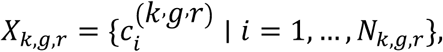

where *c_i_^(k,g,r)^* denotes the barcode of the *i*-th cell in that subset and *N*_*k*,*g*,*r*_ is the number of cells in the subset.

From each subset *X*_*k,g,r*_, CellDEEP selects *n* cells to pool and aggregates their gene read counts to construct metacells. The pooling size *n* is user-defined, with a minimum of 1. If a subset contains fewer than *n* cells, it is discarded.

### Selecting cells to pool from subset

For cell selection, we implemented two strategies: random selection and k-means clustering selection.

Assuming *n* < *N*_*k,g,r*_, for random selection we randomly select *n* cells from each *X*_*k,g,r*_can pool the cells. The *n* cells will form one metacell in further aggregation and are then removed from *X*_*k,g,r*_. Sampling is then repeated for the next *n* remaining cells until all cells in *X*_*k,g,r*_have been processed. The final metacell will contain the remaining cells.

For k-means clustering selection, the number of clusters, *K,* is set according to the intended pooling size *n* and *N*_*k,g,r*_:

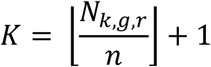

If *K* <= 2, the procedure is not applied because the subset contains too few cells for meaningful clustering (in which case a smaller *n* is required). Otherwise, *k*-means clustering is performed on the variance-standardized PCA embedding of *X*_*k,g,r*_ to obtain *K* clusters. Cells within each cluster are then merged to form one metacell per cluster.

### Aggregating gene read counts

For each pool of *n* selected cells from subset *X*_*k,g,r*_, CellDEEP aggregates gene read counts to construct a metacell.

Let *y*_*ij*_ denote the read count of gene *j* in cell *c*_*i*_.

For a pool *P* ⊂ *X*_*k,g,r*_ with ∣ *P* ∣= *n*, aggregated counts for gene *j* are computed as:

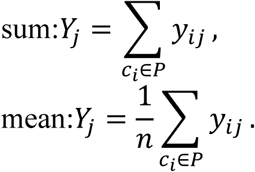

The resulting vector, *Y* = (*Y*_1_, …, *Y*_*G*_), defines the read count profile of one metacell, *G* is the number of genes. *X*_*k,g,r*_Pooling and aggregation are repeated until all cells in subset are processed.

This process is applied to all subset datasets, the resulting metacell vectors are then combined to construct a complete metacell expression matrix. Finally, a pooled Seurat object is constructed from the aggregated metacell matrix.

### Application of DE methods

Differential expression analysis in CellDEEP follows the standard Seurat^15^ workflow and is performed using the *FindMarkers* function. All parameters are configurable as in the standard Seurat pipeline, with a default *p.adj (adjusted p-value)* threshold of 0.05. For scRNAseq-based DE methods we used MAST and DESeq2, through Seurat package v. 4.3.0, with default parameters, and FindMarkers *p_val_adj* < 0.05. For pseudobulk-based methods we used DESeq2 and Limma-voom, through *pbDS()* function from Muscat^4^ Version 1.12 package (with default parameters *p_adj.loc* < 0.05; *min_cells*=1). Before DE analysis, the Seurat object is converted to the *SingleCellExperiment* object, and then the pseudobulk data matrix is generated by summing gene read counts across cells from the same group, cluster and biological replicates. The design matrix, which is used in differential expression comparison, is calculated by the combination of “*cluster_id*” and “*sample_id*”. All processing steps were carried out using the standard Muscat pipeline.

For CellDEEP, we also used MAST and DESeq2 from Seurat package v. 4.3.0 as analysis methods after pooling.

### Simulation

To evaluate the performance and utility of CellDEEP, we first tested it on simulated datasets. We used the simulation framework from two published simulation pipelines: Crowell et al^4^ and Zimmerman et al^3^. To maximize the similarity of the simulated dataset, we used the same input settings for both frameworks (indicated respectively as Muscat and Zimmerman): we generated datasets with 4000 cells, 2000 genes, 2 groups, and each group with 10 replicates, 20% DE genes. For Muscat, five reference datasets were used: the original reference dataset from Zimmerman (Alpha cells from pancreas)^3^ and Muscat^4^ (PBMC from Lupus patients obtained before and after 6h-treatment with IFN-β), CD14^+^ monocytes and CD8^+^ T cell (in healthy and severe COVID-19 patients)^16^ and TREM2^high^ macrophage cluster (from rheumatoid arthritis active patients and healthy donors)^17^. The above datasets were also used for the Zimmerman pipeline with the exclusion of the PBMC dataset from Lupus patients as it failed to meet the framework requirements.

In particular, for Muscat framework (Muscat v. 1.12.1) we simulated a total of 10 datasets. For each reference dataset, we generated two simulation settings with average log_2_ fold changes (log_2_FC) of 2 and 3, respectively.

Muscat implements multiple differential expression patterns, including changes in the mean expression (DE), changes in the proportions of cells expressing a gene (DP), changes in differential modality (DM), and changes in both proportions and modality (DB). Genes that are not subject to changes are either equivalently expressed (EE) or expressed at low and high levels in equal proportions (EP) across conditions. Full details of these simulation parameters are provided in the original Muscat publication^4^. To keep the comparability between two simulation frameworks, we set 20% DE, 0% DP, 0% DM, 0% DB, 40% EE, 40% EP for all 10 datasets.

For the Zimmerman simulation instead, we used the Murphy & Skene adaptation of the original Zimmerman framework to jointly evaluate type I and type II error^18^. Simulations were performed using the *simulate_hierarchicell* function. For each reference dataset, a non-DE matrix and a DE matrix were generated separately using identical simulation settings and then were combined into a single expression matrix. Genes from the DE matrix were treated as ground-truth DE genes, and genes from the non-DE matrix were treated as non-DE genes. The combined matrix was then converted into a Seurat object. For each reference, we simulated two datasets with linear fold change (FC) of 4 and 8 separately, to match log_2_FC used in Muscat, for a total of 8 datasets.

### Simulation analysis and result evaluation

We evaluated differential expression methods on all simulated datasets using three categories of approaches: (1) single-cell methods (DESeq2, MAST) applied directly to the original data; (2) CellDEEP with all parameter combinations (pool size, cell selection: random or k-means, aggregation: mean or sum) followed by DESeq2 or MAST; and (3) pseudobulk methods (DESeq2, limma).

Method performance was assessed using counts of true positives (TP), false positives (FP), true negatives (TN), and false negatives (FN), from which we computed precision, sensitivity, and accuracy.

To test whether performance differences between methods were statistically significant, we used a paired Wilcoxon signed-rank test (*wilcox.test* function from the R stats package, with paired = TRUE and default parameters).

### Dropout Rate

Dropout rate was defined following design of Zimmerman^3^. For each gene *g*, the dropout rate *D*_*g*_ was calculated as the proportion of cells with zero read counts among all cells.

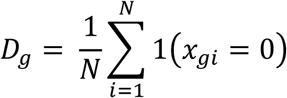

Where *N* represents the total number of cells, *x*_*gi*_ represents the read count of gene *g* in cell *i*. 1(*x*_*gi*_ = 0) is an indicator function that equals to 1 if *x*_*gi*_ = 0 and 0 otherwise.

### Published dataset test

Two published datasets are used in both simulation (as reference) and real dataset tests. To maximise similarity, we preprocess them in the way described below.

### COVID-19 PBMC

The h5ad file was downloaded from https://covid19cellatlas.org/ “COVID-19 PBMC Ncl-Cambridge-UCL” - Haniffa Lab, and transformed into a Seurat object with the *Convert* function. For the COVID-19 dataset: CD8^+^ T cells, B cell, CD14^+^ Monocytes and NK cells CD56^high^ cluster were selected for testing. Further details of datasets can be found in Supplementary Table 1. For each selected cluster, we subset them from the original COVID-19 data, DE analysis between healthy and severe COVID-19 patients was then performed. Preprocessing consists of *NormalizeData, ScaleData, RunPCA* with default parameters and 30 dimensions from Seurat package for k-means cell selection. DE analysis between healthy and severe COVID-19 patients was then performed.

### Rheumatoid arthritis

The RDS object was provided upon request from the corresponding research group^17^, TREM2^high^ and HLA^high^CLEC10A^high^ clusters were selected for downstream analysis. Data details in Supplementary Table 2. Preprocessing includes *NormalizeData, ScaleData, RunPCA* with 30 dimensions from the Seurat package for k-means cell selection. DE analysis was then performed, with comparisons between “Healthy” and “Active RA”.

### Real datasets evaluation

Two downstream analyses were performed to evaluate method performance: Null hypothesis p-value test for FP rate (FPR) and gene-ontology (GO) enrichment analysis for pathway recovery rate and signal density. We used the R package clusterProfiler v. 4.6.2 to perform GO terms enrichment analysis on identified DE genes (GO terms set as “ALL”).

### Null hypothesis p-value histogram

*Null hypothesis* data was generated by aggregating biological replicates from the same biological condition (e.g., healthy) into two separate virtual groups. For each cluster, the condition with most replicates was split in two groups, applying differential expression analysis between these two groups (Supplementary Table 3). A gene list with p-values were generated after DE analysis. Under the null hypothesis, the p-value of all genes should be uniformly distributed over [0,1], therefore a 5% false positive rate is expected. Genes with p-value < 0.05 are considered as FP results.

To evaluate method performance, the false positive rate (FPR) was calculated by dividing the number of false positive genes by the total number of genes in the corresponding null hypothesis dataset.

### Gene Ontology analysis

To assess TP detection on real data, we predefined, for each disease context, a set of GO terms expected to be enriched based on established biological knowledge. All selected GO term sets are provided in Supplementary Table 4.

For each method, we quantified TP performance using two metrics: pathway recovery rate and signal density. Let *G* denote the selected GO term list, *D* represents the set of GO terms detected by a method, and *N*_*DE*_ the number of DE genes detected by that method.

Pathway recovery rate measured the proportion of expected GO terms recovered, defined as:

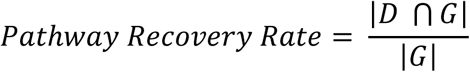

Signal density measured the number of recovered expected GO terms per detected DE gene, and thus reflecting precision in capturing biologically relevant signal rather than increased DE gene calls, defined as:

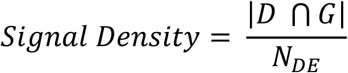

For the COVID-19 PBMC data, we focused on cell types involved in antiviral defence: antigen-presenting cells (monocytes CD14^+^, B cells) and effector cells (CD8^+^ T cells, NK cells CD56^high^) that respond to virus by activating innate immunity pathways and mechanisms. For these, we selected 91 related GO terms capturing key innate immune processes, including antiviral responses, interferon signalling, antigen processing and presentation, and inflammatory pathways.

For RA datasets, we defined two separate true positive sets: (1) for genes upregulated in healthy synovial tissue, we selected 23 GO terms related to normal tissue homeostasis; (2) for genes upregulated in active RA, we selected 46 GO terms associated with chronic inflammation, including antiviral and interferon responses, antigen presentation, and chemokine activity, consistent with established literature on RA pathogenesis^19^. For the HLA^high^ cluster, where healthy baseline biology is less clearly defined, we focused exclusively on genes upregulated in active RA, selecting 105 GO terms related to viral response, antigen presentation, and MHC-related processes.

## Results

We developed CellDEEP, an R package to improve differential-expression (DE) process by pooling single cells into metacells. CellDEEP aggregates cells within user-defined clusters into metacells. The method offers several user-configurable parameters: (1) cells can be selected for pooling either randomly within a cluster or based on k-means clustering in the embedding space (Supplementary Figure 1); (2) UMI counts for the metacell can be generated by summation (“sum”) or averaging (“mean”); and (3) the number of cells per metacell is adjustable. The resulting metacell count matrices can then be analysed using established differential expression tools such as DESeq2 or MAST.

### Exploring the best parameters set for CellDEEP

To identify optimal parameter settings, we first evaluated how count matrix generation methods (sum vs. mean) and cell selection strategies (k-means vs. random) affect performance using two established simulation frameworks: Muscat and Zimmerman (see Methods), comprising 18 simulated datasets with diverse cell types and experimental conditions.

The most substantial performance difference was observed between count aggregation methods, where, across both simulation frameworks, sum consistently outperformed mean in accuracy and sensitivity (Figure 1A-B). This effect was most pronounced in the Zimmerman simulation with MAST, where sum outperformed mean by 7 percentage points in accuracy (p≤0.08).

**Figure 1.**
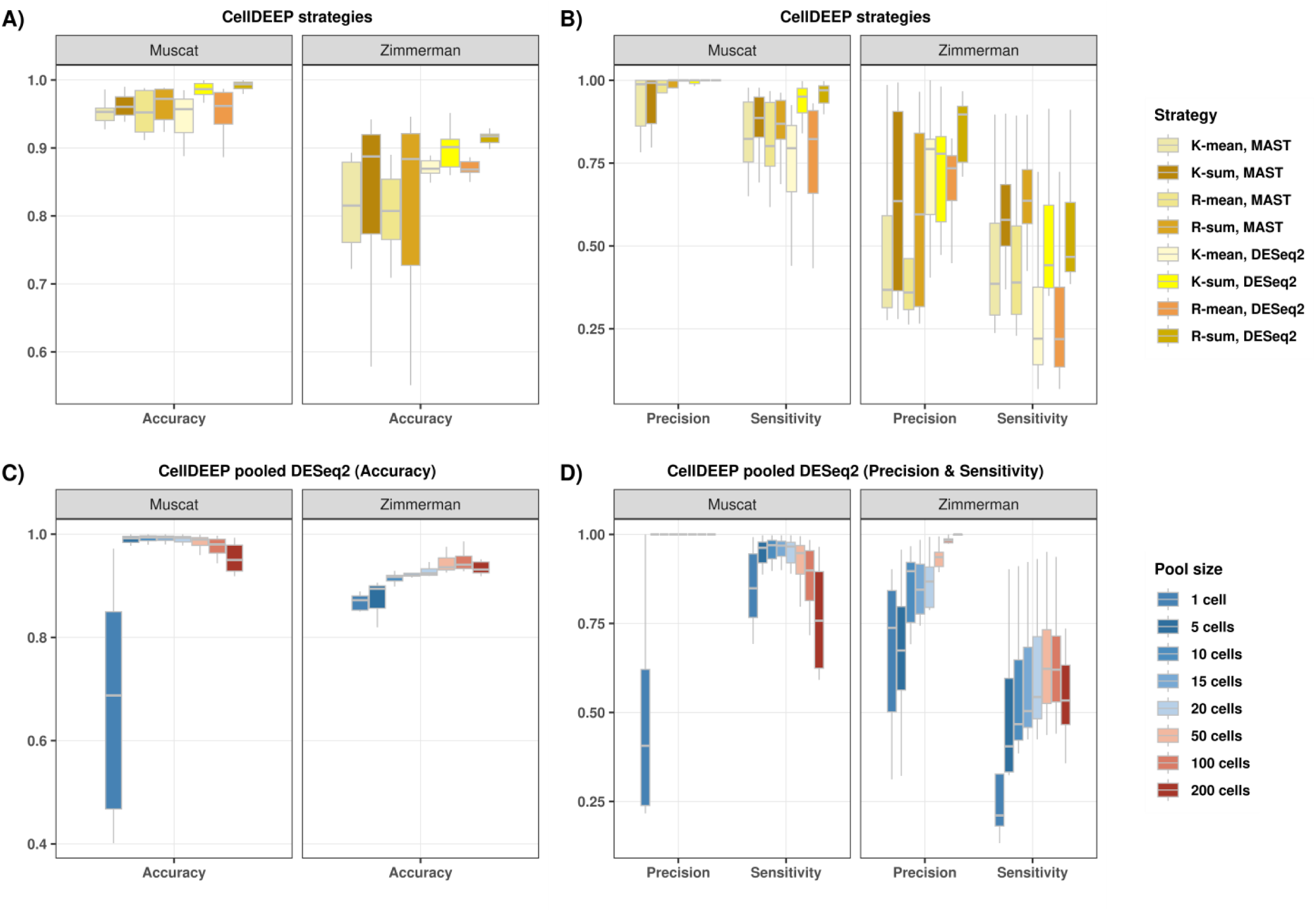
CellDEEP strategy performance comparison in simulated datasets. **A-B)** Comparison of accuracy, precision and sensitivity across both Muscat and Zimmerman simulation frameworks. K-: Cell selection by k-mean clustering; R-: Random cell selection; sum: aggregating readcounts by summing; mean: aggregating readcounts by averaging. **C-D)** Effect of metacell size on accuracy, precision and sensitivity, using optimal parameters (random selection, sum aggregation, DESeq2), in both Muscat and Zimmerman simulation framework. Metacell sizes tested: 1 (single-cell), 5, 10, 15, 20, 50, 100, and 200 cells.

In contrast, our analysis revealed that the choice of cell selection strategy (k-means vs. random) had a minimal impact on performance, with a non-significant difference in accuracy of only 1-2 percentage points (Table 1).

**Table 1.** Comparison of CellDEEP strategies for accuracy in the Zimmerman simulation. Read-count aggregation methods (Mean vs. Sum) and cell-selection approaches (K-means vs. Random) are evaluated over two DE methods. Values represent the median accuracy across 10 simulated Zimmerman datasets.

| Aggregation parameter | Cell selection parameter | Accuracy |  |
| --- | --- | --- | --- |
|  |  | MAST | DESeq2 |
| Mean | Kmean | 0.82 | 0.87 |
|  | Random | 0.81 | 0.87 |
| Sum | Kmean | 0.89 | 0.90 |
|  | Random | 0.88 | **0.92** |

The best results in terms of accuracy (0.92) were obtained by using a random selection strategy, summing the read counts and using DESeq2 (R-sum, DESeq2). These trends were consistently observed in the Muscat simulation framework as well (Supplementary Table 5).

To better understand the drivers of accuracy, we also assessed precision and sensitivity separately (Figure 1B). This revealed that sensitivity was strongly dependent on aggregation method, with sum yielding significantly higher values than mean (p < 0.03). For instance, in the Zimmerman simulation, using random selection with DESeq2, the sum aggregation (R-sum, DESeq2) achieved median precision and sensitivity of 0.90 and 0.47, respectively, whereas the mean aggregation under the same conditions (R-mean, DESeq2) yielded lower values of 0.73 and 0.22.

Based on these results, we conclude that sum aggregation consistently outperforms mean in both accuracy and sensitivity. The homogeneity of cell clusters within these simulated datasets appears to limit the potential for selection strategies to yield significant performance gains. Our analysis also highlighted a clear influence of the simulation framework itself, with Muscat yielding higher overall precision and sensitivity than Zimmerman across all parameter combinations.

### The effect of pooling cells to metacells

After establishing optimal parameters, we evaluated the effect of metacell size on detection accuracy. For each simulation we generated 4000 cells, and we varied the number of cells pooled per metacell while maintaining optimal parameters (R-sum, DESeq2). We tested extreme cases, ranging from single-cell (standard scRNA-seq) to large pools of 200 cells. While results differed slightly between simulation frameworks, they revealed consistent pattern: accuracy improved as more cells were pooled, reached an optimum, and then declined with the largest pool sizes (Figure 1C). In the Zimmerman simulations, accuracy increased incrementally from 0.87 (single-cell) to a peak of 0.94 with 100 cells per metacell, before decreasing slightly to 0.93 with 200 cells. The Muscat simulations showed a different trajectory: accuracy rose sharply from 0.75 (single-cell) to approximately 0.99 with moderate pool sizes (5–20 cells), where it remained stable, before to falling to 0.94 at 200 cells.

In both simulations, the lowest accuracy was observed with single-cell analysis (no pooling), corresponding to conventional scRNA-seq methods (Figure 1C). Accuracy improved with pooling, reaching an optimum at 20 cells for Muscat and 100 cells for Zimmerman, before declining with the largest pool sizes.

To investigate this further, we complemented our accuracy analysis with precision and sensitivity, revealing the drivers of this pattern (Figure 1D), as both were strongly affected by pool size. In Zimmerman simulations, precision increased progressively, gaining approximately 20 percentage points from pools of 5 to 200 cells. In Muscat simulations, precision rose sharply from 0.75 (single-cell) to a near-perfect 1.0 with any level of pooling. In general, by pooling sufficient cells, we can achieve a precision of 1 or close to it.

Sensitivity instead, followed a different trajectory. In Zimmerman simulations sensitivity peaked at 0.62 with 50 cells, followed by a decline to 0.53 at 200 cells. Muscat simulations followed a similar pattern of increase and then decrease, though the specific values and peak location differed. This decline in sensitivity with the largest pools explains the corresponding drop in overall accuracy.

In summary, both frameworks exhibited dramatic improvements when metacells were used compared to single-cell analysis. These results demonstrate that metacell pooling substantially enhances scRNA-seq analysis for both simulation frameworks. The gains in precision and sensitivity, at least at intermediate pool sizes for Muscat, highlight the utility of metacells in balancing statistical power and resolution. This approach mitigates noise inherent in single-cell data while avoiding the oversimplification of full-cluster aggregation, where sensitivity is lost. On the other hand, in practice, the maximum feasible pool size is constrained by the minimum number of cells per cluster available across samples to be pooled. Setting the pool size too high risks excluding samples with lower cell counts, which could reduce statistical power. Therefore, while our simulations show that pooling improves accuracy, the choice of pool size for a given study should balance these accuracy gains against the risk of sample loss.

### Comparative performance of CellDEEP against scRNA-seq and pseudobulk methods

After exploring how to parameterise CellDEEP, we next evaluated the performance of CellDEEP against conventional single-cell (DESeq2, MAST) and pseudobulk approaches (DESeq2, limma). While performance trends were consistent across simulation frameworks, specific outcomes exhibited framework-dependent variations. Based on the previous section, we decided to use 20 as the number of cells to pool. As shown previously in the simulations generated by the Zimmerman framework, CellDEEP (R-sum, DESeq2) demonstrated a 5-percentage-point increase in accuracy over standard scRNA-seq DESeq2. Furthermore, CellDEEP outperformed MAST by 10 percentage points. A similar trend was observed in the simulations generated by Muscat (Table 2).

**Table 2.**
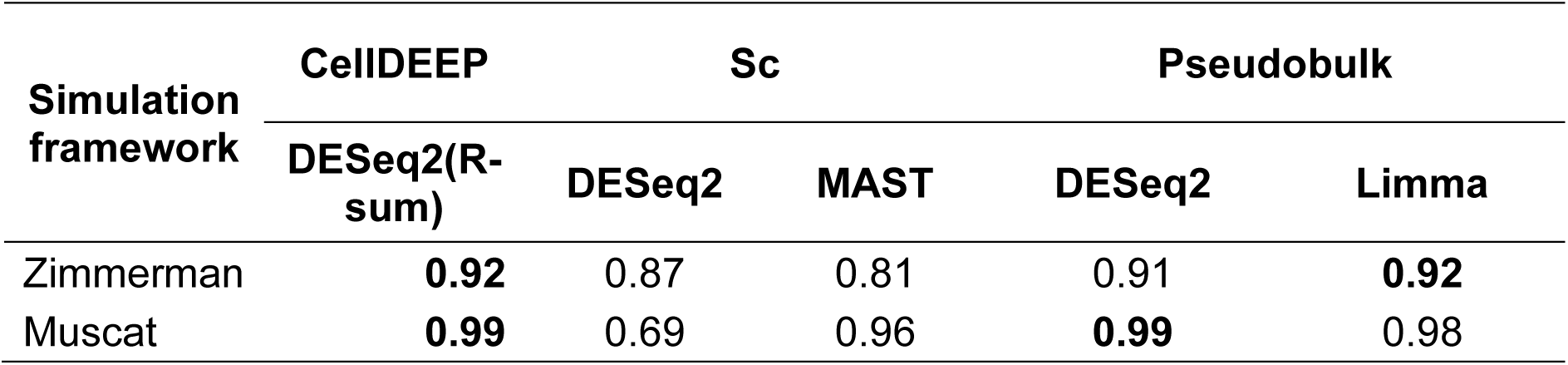
Accuracy (median) of differential expression approaches across two simulation frameworks. Comparing CellDEEP (Random–Sum–DESeq2), scRNA-seq (DESeq2, MAST), and Pseudobulk (DESeq2, Limma).

In comparison with pseudobulk methods, CellDEEP achieved very similar results in terms of overall accuracy (Table 2, Figure 2A). However, its performance in precision and sensitivity was simulation framework dependent. While CellDEEP matched pseudobulk accuracy in Muscat simulations, results from the Zimmerman framework revealed distinct trade-offs (Supplementary Figure 2): CellDEEP maintained higher sensitivity (0.61) compared to both pseudobulk DESeq2 and limma (0.54).

**Figure 2.**
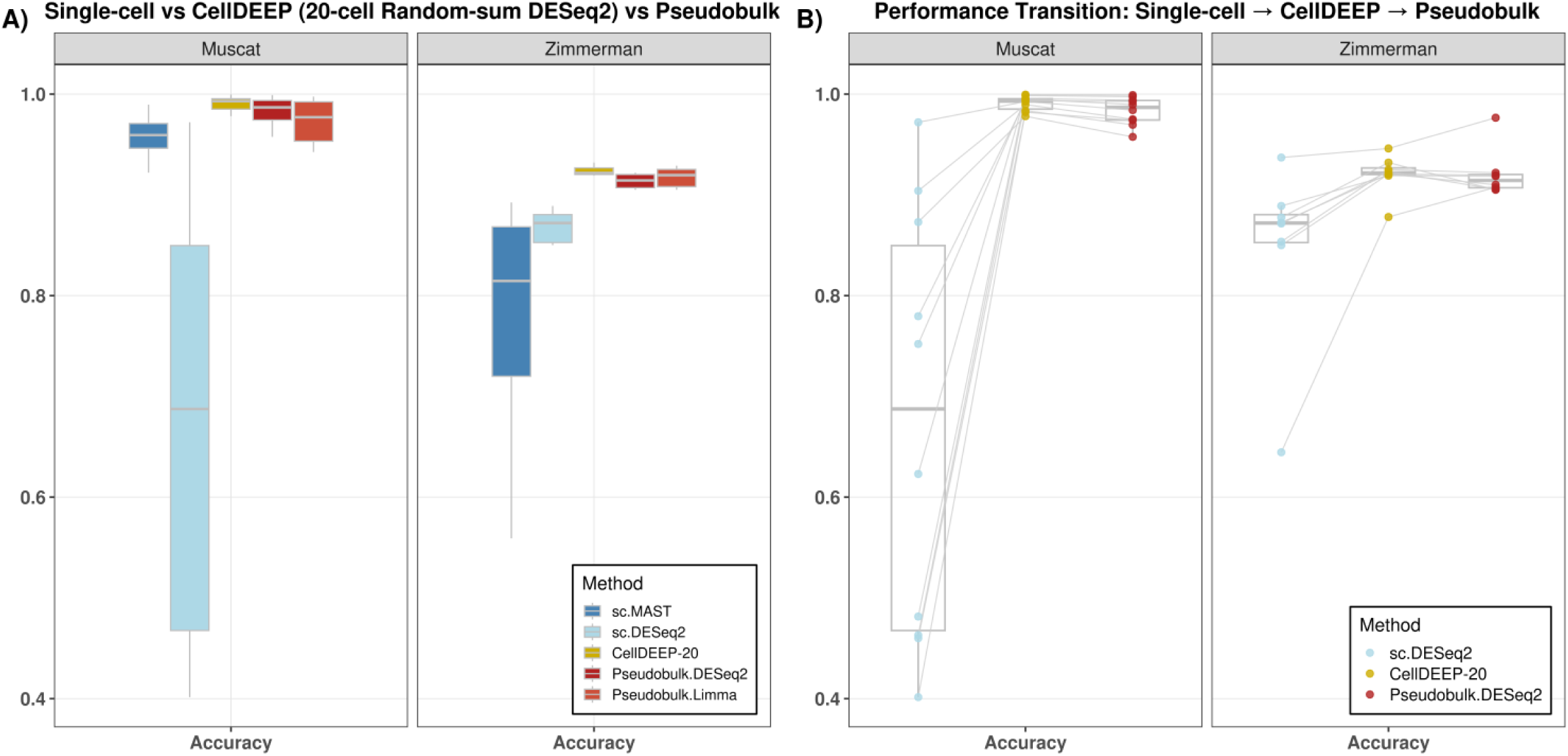
CellDEEP performance evaluation with scRNA-based and pseudobulk-based methods in simulated datasets. **A)** Accuracy of differential expression approaches across two simulation frameworks, comparing CellDEEP (Random–Sum–DESeq2), scRNA-seq (DESeq2, MAST), and Pseudobulk (DESeq2, Limma). **B)** Accuracy changes across same dataset (connected by lines) with different DE analysis methods, in two simulation frameworks comparing CellDEEP (Random–Sum–DESeq2), scRNA-seq (DESeq2), and Pseudobulk (DESeq2). Each line represents a dataset.

Conversely, CellDEEP showed moderately lower precision (0.83) than limma (0.91) and pseudobulk DESeq2 (0.88).

Our simulation results demonstrate that CellDEEP reliably and substantially outperforms standard single-cell analysis methods, likely due to a reduction of technical noise and dropout effects (Supplementary Figure 3). In comparisons with pseudobulk methods, however, the performance difference was smaller and more variable. CellDEEP achieved higher accuracy than pseudobulk in 14 out of 18 simulated datasets (Figure 2B). This variability, even within controlled simulations, highlights that differential expression analysis is inherently dataset dependent.

These findings show that CellDEEP outperforms scRNA-seq and is as good as pseudobulk on simulated data. However, simulation may favour pseudobulk methods, as it cannot capture real noise. Therefore, evaluation using real-world data is necessary to provide a more realistic assessment.

### CellDEEP outperforms both pseudobulk-based and scRNAseq-based methods on real datasets

To evaluate CellDEEP’s performance on real biological data, we analysed multiple immune cell clusters from two public datasets: a COVID-19 PBMC dataset and a RA synovial tissue dataset.

Our evaluation strategy was based on two principles: (1) assessing false positives by testing under the null hypothesis, and (2) evaluating true positive detection by examining expected GO terms and their associated genes from well-established biological processes. We used two papers well-established in the field: for COVID-19, we analysed antigen-presenting cells (CD14^+^ monocytes, B cells) and antiviral effector cells (plasmacytoid DCs, CD8^+^ T cells, NK cells CD56^high^) from healthy vs. severe patient samples; for rheumatoid arthritis (RA), we analysed synovial macrophages, which play a key role in immune response and tissue homeostasis in the RA synovial membrane, and compared them with healthy controls^17,20,21^. For these analyses, we used a metacell pool size of 10 cells to ensure that sufficient metacell replicates could be generated from all clusters, including those with limited per-sample cell counts (e.g., NK cells CD56^high^).

### False positive rate

To estimate the false positive rate (FPR), we performed negative control comparisons where no differential expression should exist^22^. This was achieved by splitting the replicate samples from a single biological condition (e.g., healthy donors) into two artificial groups. Under these conditions, genes with an adjusted p-value < 0.05 after DE were classified as false positives.

Overall, MAST exhibited the poorest performance across single-cell (FPR 0.3–0.6), compared to MAST with CellDEEP (R-sum) (0.12–0.55), and CellDEEP (R-mean) (0.08–0.16) configurations (Figure 3A). In our experiment and tested dataset, MAST failed to control false positives well compared to other scRNA-seq methods. Both scRNA-seq DESeq2 and pseudobulk limma also exceeded the 0.05 false positive threshold. The two best-performing tools were pseudobulk DESeq2 (FPR ≤ 0.02) and CellDEEP (R-mean, DESeq2) (FPR ≤ 0.03), both of which maintained false discovery rates below the 0.05 cutoff. Results were consistent across individual datasets, with CD8+ T cells presenting the greatest challenge (FPR range 0.01–0.6) and B cells the least (FPR range 0.01–0.3).

**Figure 3.**
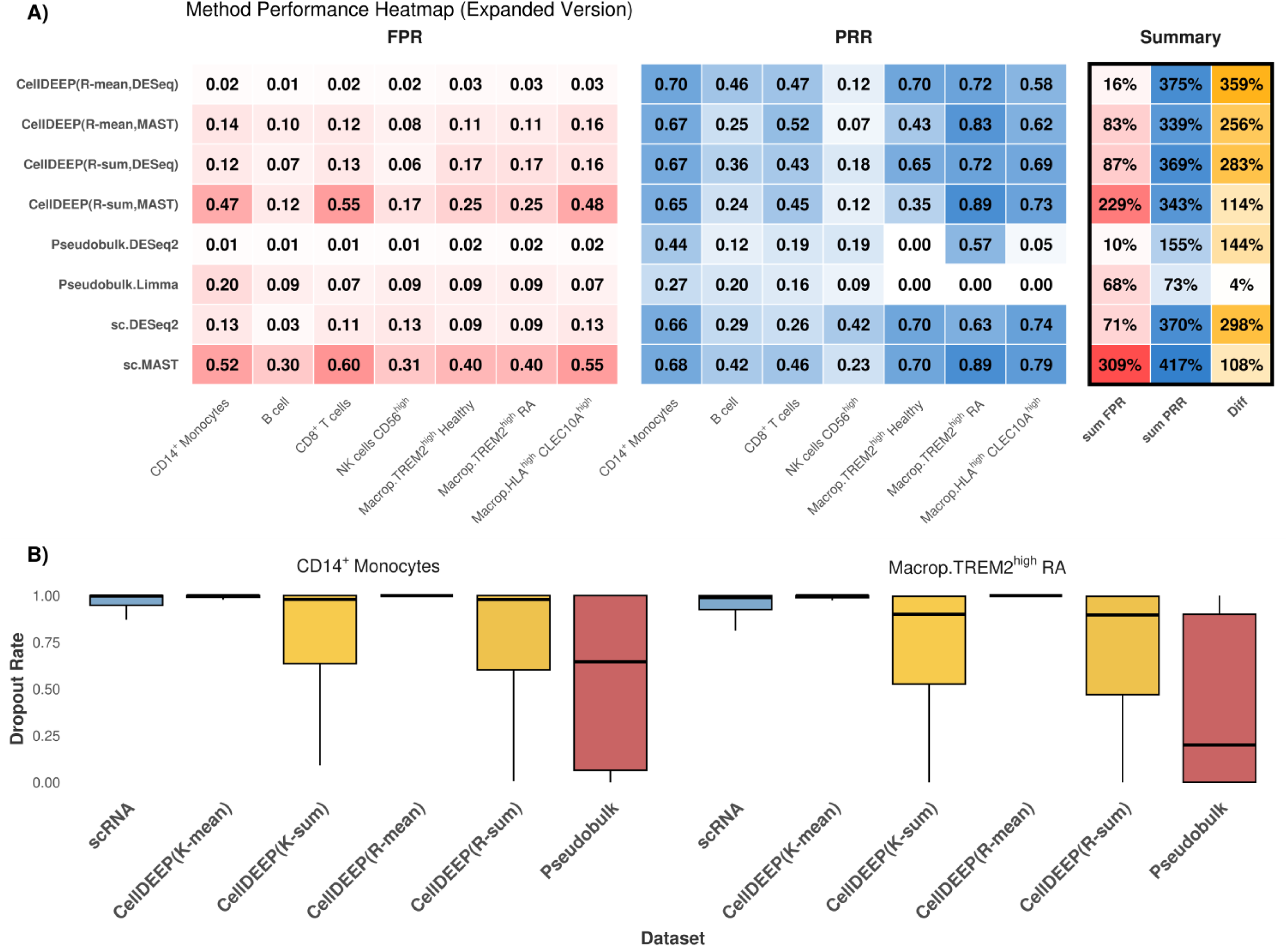
Method evaluation on published datasets: false positive rate and pathway recovery rate. **A)** Performance summary across all datasets. For each method, we calculated: false positive rate (FPR) from negative control comparisons (same-condition replicate splits); pathway recovery rate (PRR) as the proportion of curated true positive GO terms detected. Sum FPR and Sum PRR represent totals across all datasets (percentages). Difference (Sum PRR − Sum FPR) indicates the net balance between true positive detection and false positive control. Methods evaluated: scRNA-seq (MAST, DESeq2); pseudobulk (DESeq2, Limma); CellDEEP (random selection, 10 cells per metacell) with sum/mean aggregation and DESeq2/Limma. **B**) Gene dropout comparison for CD14^+^ monocytes (COVID-19) and TREM2^high^ macrophages (RA). For each gene, dropout rate was calculated as the proportion of cells with 0 read counts. Methods evaluated i: original scRNA-seq data; CellDEEP (k-means or random selection, sum or mean aggregation, 10 cells per metacell); and pseudobulk

To better understand these differences, we examined the negative control comparisons in two clusters in greater detail by visualising the p-value distributions (Supplementary Figures 5 and 6). In an ideal negative control experiment, only 5% of p-values should fall below 0.05, and their distribution should be uniform between 0 and 1. In the CD14+ monocyte cluster from the COVID-19 dataset, single-cell MAST again performed poorly, identifying 8,129 false positives in the negative control comparison (Supplementary Figure 5). While CellDEEP (R-sum) only partially mitigated this issue, reducing false positives by approximately 10%, a different CellDEEP configuration emerged as a top performer. Across all real datasets and cell types tested, CellDEEP (R-mean, DESeq2) consistently achieved the lowest false positive rates. Its performance was superior to all scRNA-seq-based methods and comparable to standard pseudobulk methods in controlling false positives (Figure 3A). These results were mirrored in the RA TREM2^High^ cluster.

Furthermore, it was unexpected that the R-mean (CellDEEP) controlled FP better than the R-sum (CellDEEP), unlike in the simulation datasets. Our hypothesis was that in real single cell data, “noise” was higher when summing the counts to form metacells and was averaged out in the mean. Hence, we compared the dropout rate of the cells for the different methods in the CD14^+^ and TREM2^High^ datasets (Figure 3B). As expected, the scRNA-Seq shows high, and the pseudobulk low, dropout rates. CellDEEP with k-means selection and sum aggregation (K-sum) behaves similarly to pseudobulk, although the median is higher compared to pseudobulk. In contrast, the values of CellDEEP with k-means selection and mean aggregation read count (K-mean) approach 1.

This effect is caused by rounding during averaging and raw count generation. When the overall read count is very low (e.g., a mean of 0.3), rounding converts the value to 0. As a result, some lowly expressed genes appear to lose signal. However, such genes with low counts are often dominated by technical noise rather than true biological signal^23^. Therefore, rounding primarily removes background noise instead of meaningful expression. In the simulations (Supplementary Figure 3), we can see the similar trend. However, the muscat dropout rates have less spread, and the Zimmerman values are similar to the real dropout rates.

### True positive

As the biological processes underlying RA and COVID-19 are well characterized, established immune pathways provide a qualitative benchmark for expected differential expression patterns. To assess our ability to detect biologically meaningful signal, we examined differentially expressed genes within canonical immune pathways known to be relevant to each condition using two complementary metrics.

First, we calculated pathway recovery rate (PRR) (Figure 3A; Supplementary Figure 7A) as the proportion of expected pathways that each method successfully detected from our curated list of true positive GO terms. This metric reveals how comprehensively a method captures known biology. Here, pseudobulk methods showed a critical limitation: they lacked power, as shown by others, recovering substantially fewer expected DEG/GO than single-cell methods or CellDEEP.

The only cluster that performs similarly is the NK cells, which has a very limited number of cells (Supplementary Table 1). The traditional single cell methods are generally close to the CellDEEP results, ranging from 0.5-0.7 in most cases, however, as noted before, many false positive results were observed for scRNAseq-based methods.

Second, we calculated signal density (Figure 4) as the proportion of each method’s called DE genes that fell within expected pathways. This metric distinguishes methods that genuinely detect relevant biology from those that simply call more genes by chance. We explored the CD14^+^ monocytes in more detail, where pseudobulk-based methods recovered roughly half the number of pathway-associated true positive genes compared to single-cell methods or CellDEEP (Figure 4A). While all methods identified core pathways such as “defense response to virus”, scRNA-seq and CellDEEP methods additionally recovered a broader set of immune processes involved in the antiviral response. These included additional immune processes central to antiviral defence (“regulation of innate immune response”, “positive regulation of cytokine production involved in inflammatory response”) as well as key effector mechanisms (“antigen processing and presentation via MHC class I”, “NF-kappaB binding”) (Supplementary Table 4). A similar pattern was observed for TREM2^high^ macrophages. Notably, pseudobulk DESeq2 showed an artificially high signal density in this cluster (Figure 4B), which was driven by its extremely conservative gene calls, as reflected in its low pathway recovery rate for the same cluster (Figure 3A).

**Figure 4.**
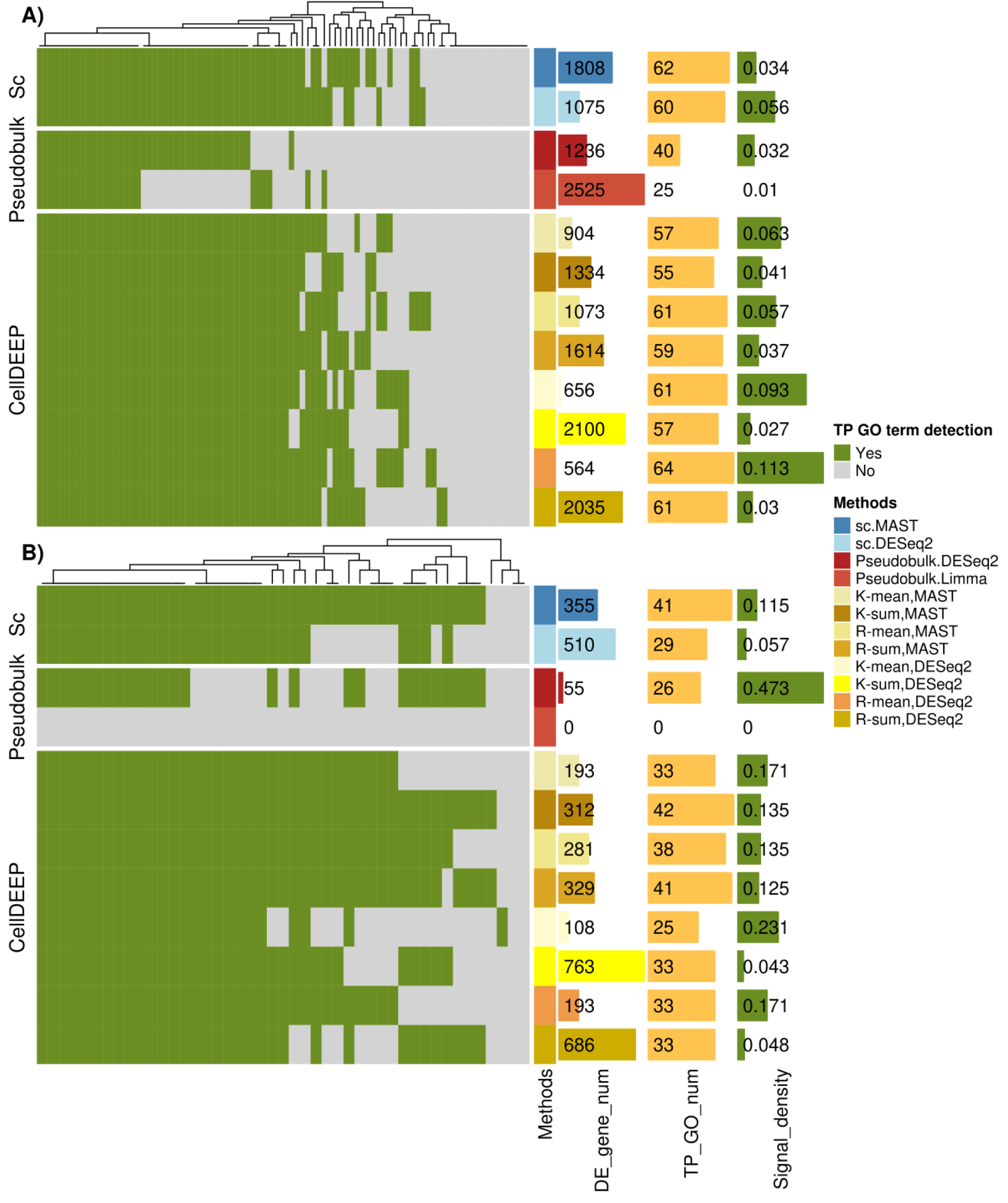
Benchmarking DE methods by true positive GO term detection. *A)* COVID19 CD14^+^ Monocytes and **B)** RA TREM2^high^ Macrophage. For each method, bars include: total number of DE genes called (coloured by method); number of true positive (TP) GO terms detected (light yellow); and signal density (green), calculated as the number of TP GO terms detected divided by total DE gene count. Methods compared: scRNA-seq (MAST, DESeq2), pseudobulk (DESeq2, Limma), and CellDEEP with various parameter combinations (selection: random or k-means; aggregation: sum or mean; downstream DE tool: DESeq2 or Limma; 10 cells per metacell).

Across all real datasets, CellDEEP methods were among the top performers by both metrics. However, the sum aggregation methods consistently generated more false positives (Supplementary Figure 7), suggesting that mean aggregation provides better specificity in real data.

Overall, CellDEEP achieved the best balance between true positive detection and false positive control. By aggregating cells into replicated metacells, it preserves sufficient biological heterogeneity to match the sensitivity of single-cell methods while stabilising expression estimates to approach the specificity of pseudobulk approaches.

## Discussion

Although single-cell transcriptomics has substantially advanced our understanding of cellular function, the robust detection of differentially expressed genes (DEGs) remains an open challenge. Previous studies have reported that single-cell differential expression methods tend to overpredict DEGs, whereas pseudobulk approaches, while better controlling false positives, may suffer from reduced power^3^. Several reports have suggested that pseudobulk represents a preferable strategy due to improved control of false discovery rates. However, these conclusions have largely been based on simulated data and benchmark where pseudobulk has been treated as gold standard, as establishing a reliable ground truth set in real scRNA-seq datasets remains inherently difficult.

First, we developed CellDEEP (Cell Differential Expression by Pooling), an approach designed to combine the strengths of single-cell and pseudobulk frameworks. To address the limitation of real data truth sets, we introduce an alternative strategy for performance assessment that leverages real datasets to approximate ground truth. Our results demonstrate that no single method consistently outperforms all others.

Conventional single-cell methods identify a large number of candidate genes but exhibit elevated false positive rates, whereas pseudobulk approaches control false positives better, at the cost of reduced sensitivity. We acknowledge that the performance of existing methods could potentially be improved through more extensive parameter tuning, for example, incorporating latent variables in MAST or including metadata covariates in pseudobulk models. However, such optimisations are equally applicable to CellDEEP and do not alter the relative performance observed in our benchmarks.

Importantly, our tool achieves a more balanced trade-off between these two extremes. Across two simulation frameworks, pseudobulk approaches showed higher overall accuracy but lower sensitivity than CellDEEP. As illustrated in Figure 1C-D, excessive pooling of cells reduces statistical power across methods. While the precise mechanistic explanation remains unclear, it is evident that simulation frameworks do not fully capture the complexity and variability inherent to real biological data^24,25^. This raises a fundamental question: how can a meaningful truth set be established using real datasets?

To address this, we propose two complementary strategies for approximating ground truth. First, we analysed large single-cell datasets from COVID-19 and RA by comparing samples within the same condition, thereby generating two biologically equivalent groups. In this setting, no true differential expression exists, as the comparison is constructed under the null hypothesis. Examination of p-value distributions (Supplementary Figures 4–5) revealed that single-cell methods produced a substantial excess of low p-values, consistent with inflated false positives. Pseudobulk methods displayed a more uniform p-value distribution, whereas CellDEEP exhibited intermediate behaviour.

To estimate true positive rates, we adopted a biologically informed strategy. For each disease, we identified Gene Ontology terms known to be relevant to disease pathology and defined genes within these categories as true positives. Under this framework, CellDEEP demonstrated strong performance, comparable to single-cell methods. Consistent with previous reports, pseudobulk approaches showed reduced sensitivity; however, this reduction was more pronounced in real datasets than in simulations.

Although CellDEEP performs robustly overall, performance varies across datasets. We did not observe a consistent pattern indicating that a specific CellDEEP implementation was universally optimal. For example, it remains unclear whether aggregation strategies such as R-sum or R-mean perform better under specific data-generating scenarios. Therefore, rather than advocating for a single superior method, we recommend the complementary use of multiple approaches, with careful consideration of their respective strengths and limitations. The principal advantage of CellDEEP lies in its balanced control of false positives and true positives.

In conclusion, we present CellDEEP, a novel framework for differential expression analysis that achieves a favourable balance between sensitivity and specificity.

Importantly, our work also introduces an alternative evaluation strategy based on real datasets rather than simulations alone, which often fail to capture the full complexity of biological variability. We suggest that CellDEEP be used alongside established methods to assess the robustness of findings and to better characterise false and true positive rates in single-cell transcriptomic analyses.

## Data Availability

The CellDEEP package is available through GitHub for download and use: https://github.com/sii-scRNA-Seq/CellDEEP

Code for all analysis presented in the paper is available through Github: https://github.com/YiyiChen0328/CellDEEP_analysis

Simulated data used during analysis are available on Zenodo: https://zenodo.org/records/18863779

## Acknowledgments

Portions of this manuscript were edited for language and clarity using a language model.

## Author contribution statement

Y.C. performed formal analysis, developed software, and generated visualizations. Y.C., T.K., R.F.L., and O.M.H. contributed to methodology. T.K., R.F.L., O.M.H., A.M. and D.S. contributed to software development. D.S. and T.D.O. conceived the study and supervised the project. D.S. wrote the original draft of the manuscript. All authors reviewed and edited the manuscript.

---

**Supplementary Figure 1.**
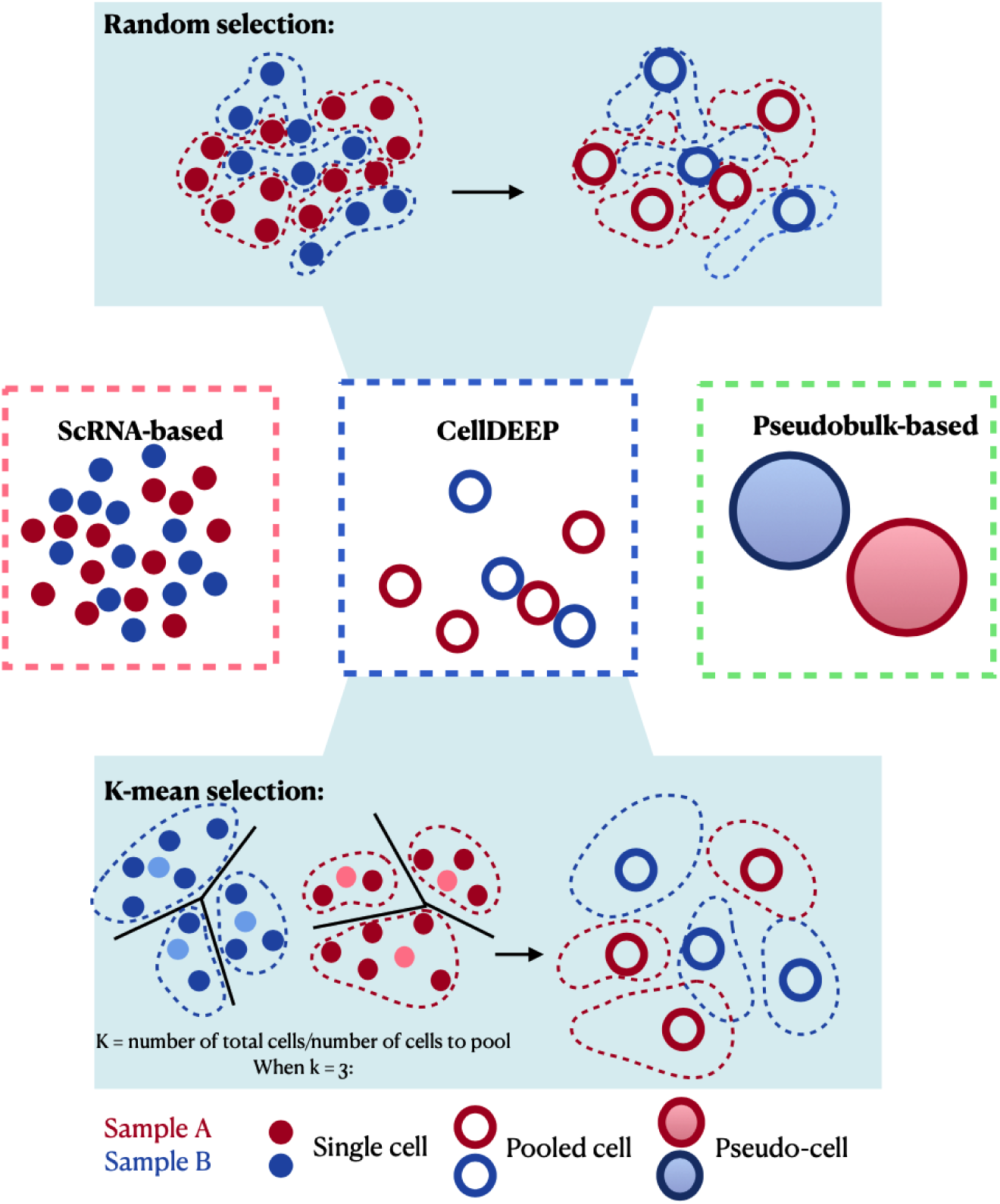
Overview of CellDEEP. Unlike scRNA-seq (which analyses individual cells) or pseudobulk approaches (which aggregate all cells in a cluster), CellDEEP aggregates individual cells into metacells using two alternative strategies: random pooling (top) or k-means clustering (bottom).

**Supplementary Figure 2.**
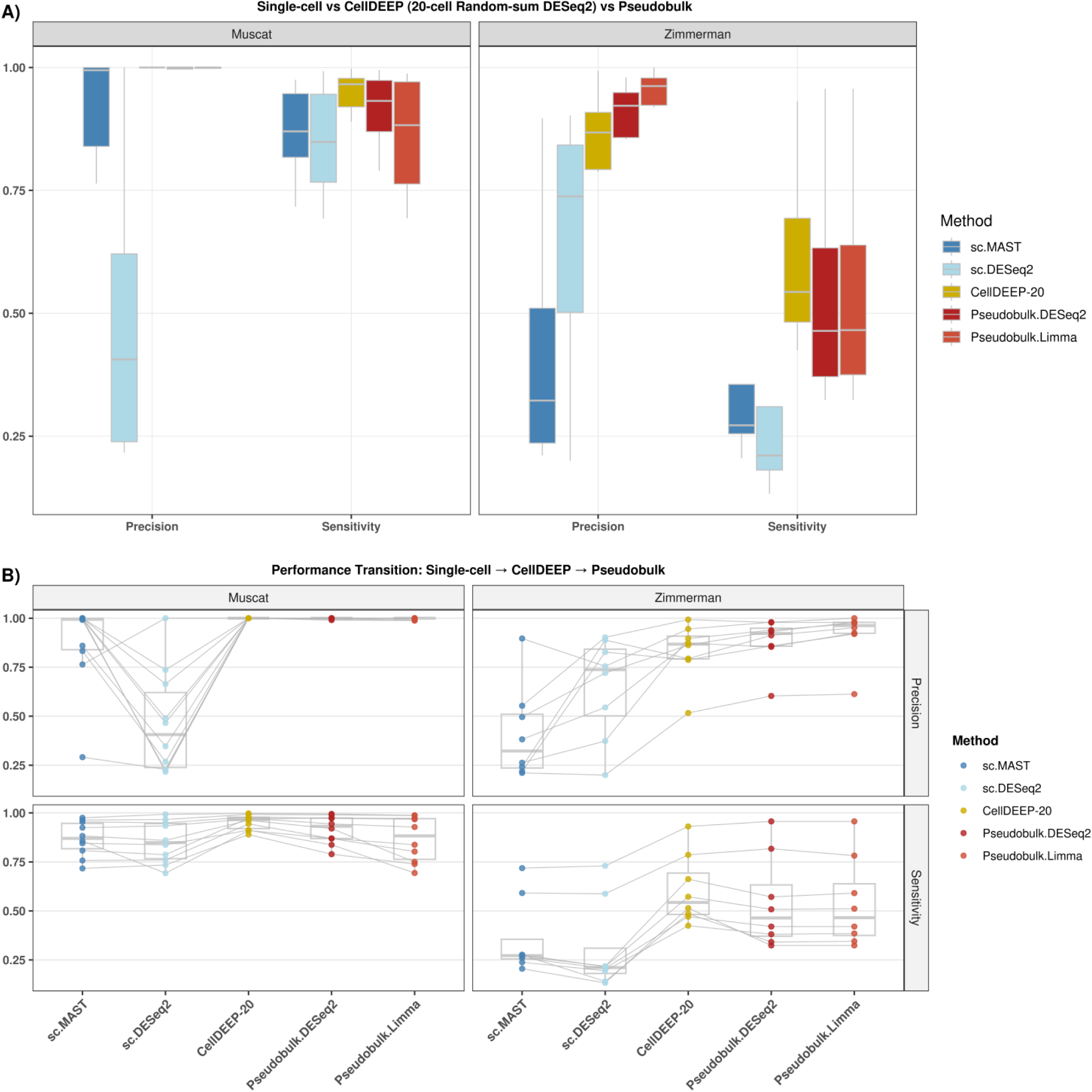
Benchmarking CellDEEP against scRNA and pseudobulk methods in simulation data. **A)** Precision and sensitivity across Muscat and Zimmerman simulation frameworks for CellDEEP (random selection, sum aggregation, 20 cells per metacell, DESeq2), scRNA-seq (DESeq2), and pseudobulk (DESeq2). **B)** Precision and sensitivity across the same datasets connected by lines, comparing CellDEEP (Random–Sum–DESeq2), scRNA-seq (DESeq2), and pseudobulk (DESeq2) in both simulation frameworks. Each line represents a shared dataset.

**Supplementary Figure 3.**
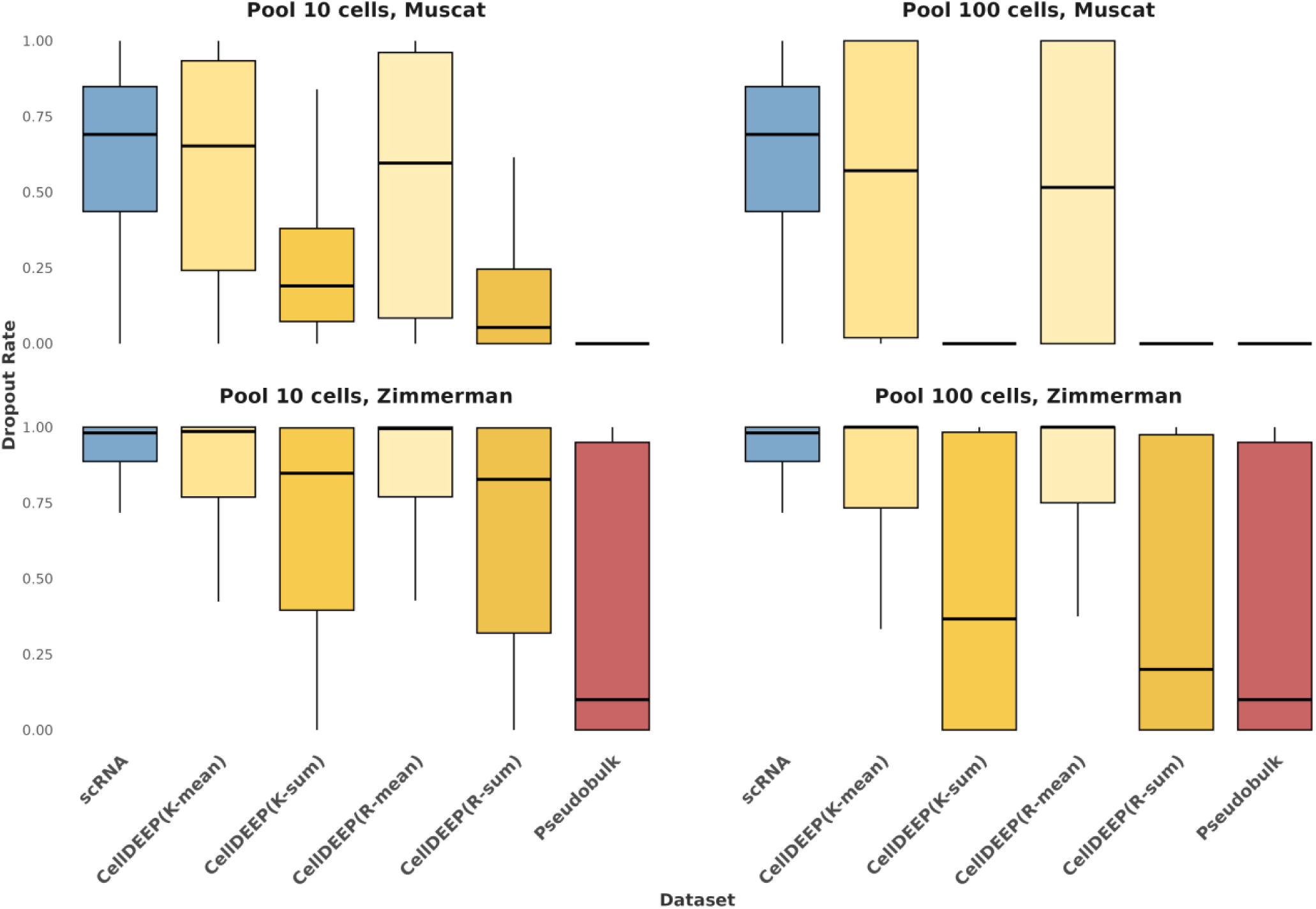
Gene dropout comparison across simulated datasets. For each gene, dropout rate was calculated as the proportion of cells with 0 read count. Results are shown separately for Zimmerman and Muscat simulation frameworks and for metacell sizes of 10 and 100 cells. Methods compared: original scRNA-seq data; CellDEEP (k-means or random selection, sum or mean aggregation, 10 or 100 cells per metacell); and pseudobulk.

**Supplementary Figure 4.**
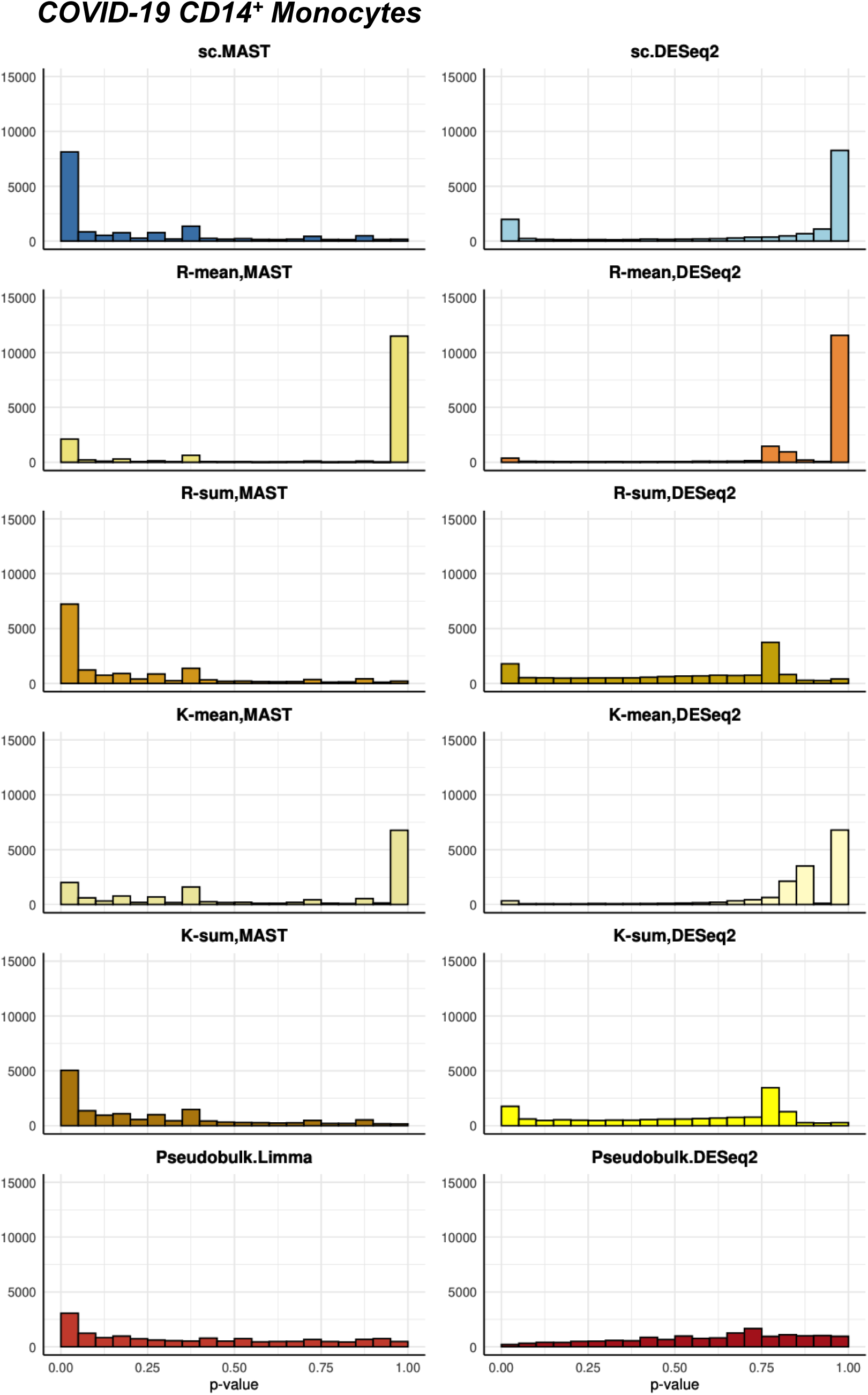
P-value distributions under null hypothesis comparisons in COVID-19 CD14^+^ Monocytes. Histograms show p-values from negative control comparisons (same-condition replicate splits) for each DE method. An ideal null distribution should be uniform with no enrichment of low p-values. Methods evaluated: scRNA-seq (MAST, DESeq2); pseudobulk (DESeq2, Limma); CellDEEP (random or k-means selection, sum or mean aggregation, 10 cells per metacell) with DESeq2 or MAST.

**Supplementary Figure 5.**
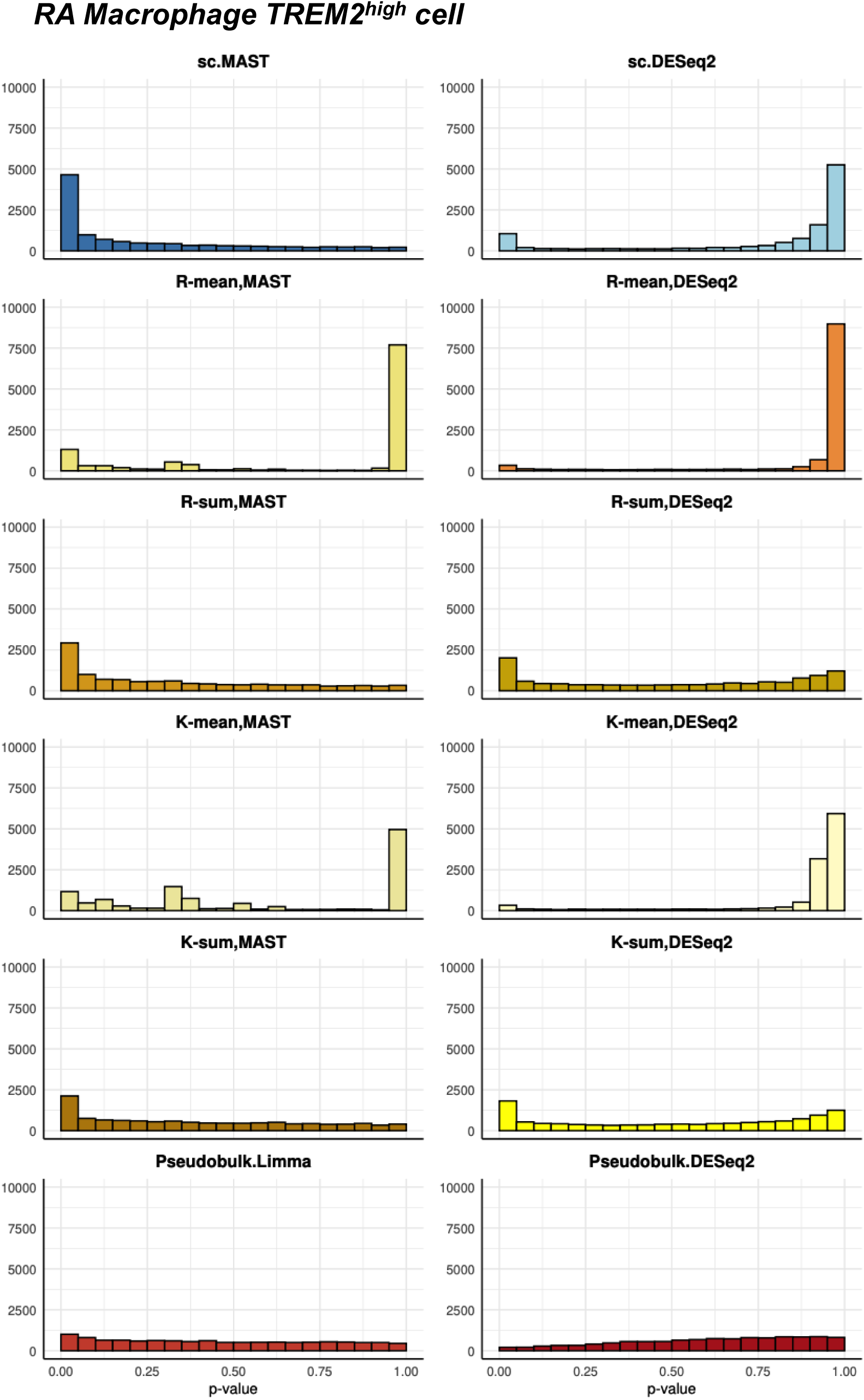
P-value distributions under null hypothesis comparisons in RA TREM2^high^ macrophages. Histograms show p-values from negative control comparisons (same-condition replicate splits) for each DE method. An ideal null distribution should be uniform with no enrichment of low p-values. Methods evaluated: scRNA-seq (MAST, DESeq2); pseudobulk (DESeq2, Limma); CellDEEP (random or k-means selection, sum or mean aggregation, 10 cells per metacell) with DESeq2 or MAST.

**Supplementary Figure 7.**
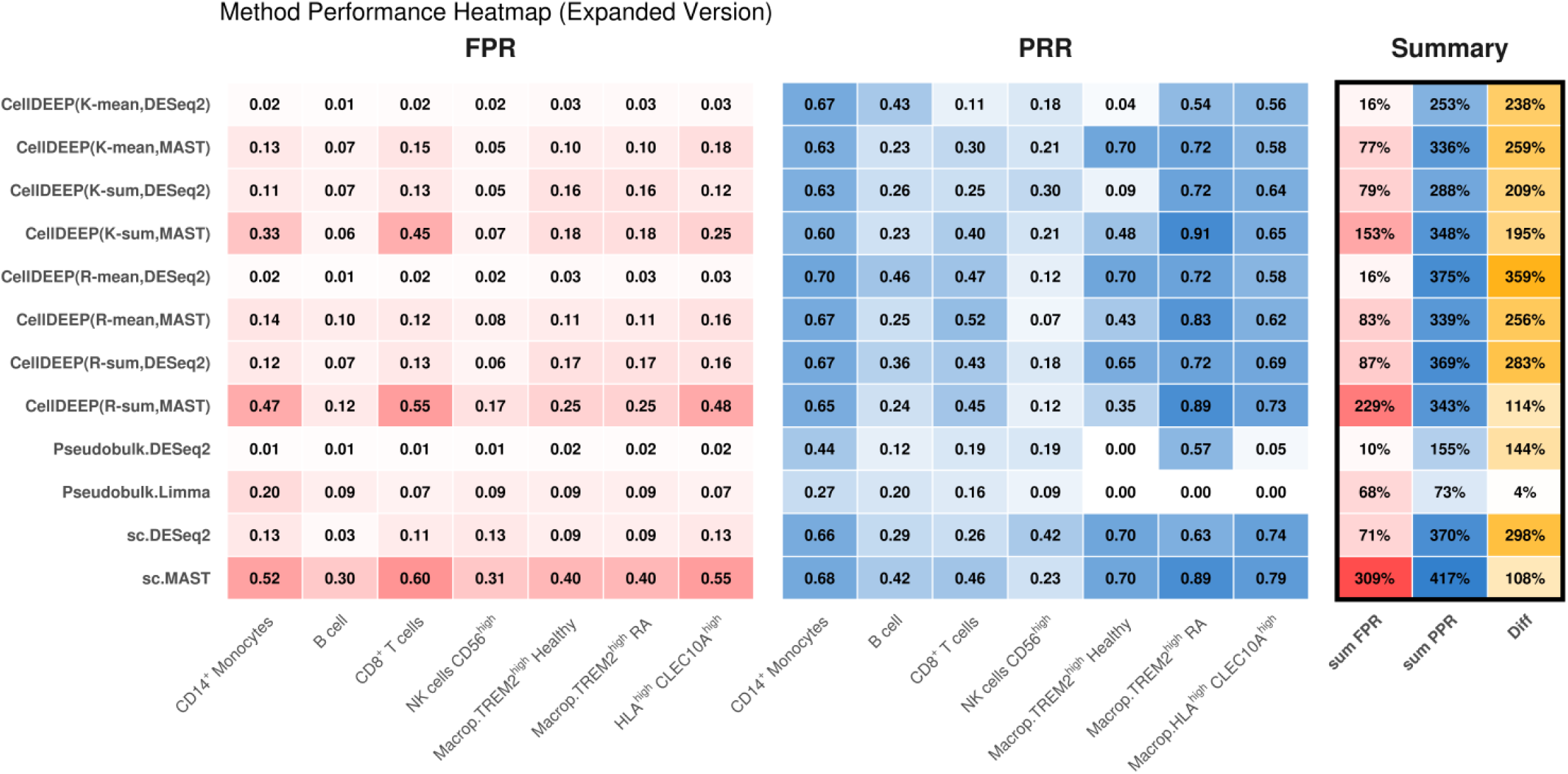
Full method evaluation on published datasets: false positive rate and pathway recovery rate. For each method, we calculated false positive rate (FPR) from negative control comparisons and pathway recovery rate (PRR) from GO enrichment analysis. Sum FPR and Sum PRR represent totals across all datasets (percentages). Difference (Sum PRR − Sum FPR) indicates the net balance between true positive detection and false positive control. Methods evaluated: scRNA-seq (MAST, DESeq2); pseudobulk (DESeq2, Limma); CellDEEP (random or k-means selection, sum or mean aggregation, 10 cells per metacell) with DESeq2 or Limma.

**Supplementary Table 1.** COVID-19 dataset summary. The replicate number and statistical summaries of cell distribution in the COVID-19 PBMC data.

| Cell type | Group | Replicant count | Total cells | Mean cells per replicate | SD cells per replicate |
| --- | --- | --- | --- | --- | --- |
| COVID-19 B cell | Healthy | 24 | 3727 | 155.29 | 161.48 |
|  | Severe | 7 | 2516 | 359.43 | 302.41 |
| COVID-19 T cells CD8<sup>+</sup> | Healthy | 21 | 9385 | 446.90 | 328.35 |
|  | Severe | 7 | 3013 | 430.43 | 503.67 |
| COVID-19 pDC | Healthy | 22 | 343 | 15.59 | 19.03 |
|  | Severe | 7 | 251 | 35.86 | 67.26 |
| COVID-19 NK CD56<sup>high</sup> cells | Healthy | 24 | 1099 | 45.79 | 37.44 |
|  | Severe | 7 | 342 | 48.86 | 26.34 |
| COVID-19 monocytes CD14<sup>+</sup> | Healthy | 24 | 5162 | 215.08 | 250.09 |
|  | Severe | 7 | 3706 | 529.43 | 514.66 |

**Supplementary Table 2.** Rheumatoid arthritis dataset summary. The replicate number and summary statistics for cell-distribution in the RA data.

| Cell Type | Group | Replicate<br>s Number | Total<br>Cells | Mean Cells Per<br>Replicate | SD Cells Per<br>Replicate |
| --- | --- | --- | --- | --- | --- |
| TREM2<sup>high</sup> | Healthy | 4 | 1775 | 443.75 | 229.54 |
|  | Active RA | 6 | 860 | 143.33 | 78.44 |
| HLA<sup>high</sup> | Healthy | 4 | 893 | 223.25 | 138.33 |
|  | Active RA | 6 | 1437 | 239.5 | 96.55 |

**Supplementary Table 3.** Null hypothesis tests summary. The table lists the sample IDs used for the p-value null hypothesis tests. For each cluster, samples from the same biological group were randomly assigned to Group 1 and Group 2, with 5 samples in each group for the COVID-19 dataset and 3 samples in each group for the RA dataset, to construct the null hypothesis test dataset.

| Cluster | Group | Samples |
| --- | --- | --- |
| B cell | 1 | MH8919333, MH8919332, MH8919227, MH8919179, MH8919177 |
|  | 2 | MH8919283, MH8919178, MH8919226, MH8919282, MH8919176 |
| CD8<sup>+</sup> T cells | 1 | MH9179822, BGCV12_CV0178, BGCV08_CV0915, MH8919179, MH8919332 |
|  | 2 | BGCV06_CV0178, newcastle65, BGCV14_CV0940, BGCV10_CV0939, BGCV05_CV0929 |
| NK cells CD56<sup>high</sup> | 1 | MH8919333, MH8919332, MH8919227, MH8919179, MH8919177 |
|  | 2 | MH8919283, MH8919178, MH8919226, MH8919282, MH8919176 |
| CD14<sup>+</sup> Monocytes | 1 | MH8919333, MH8919332, MH8919227, MH8919179, MH8919177 |
|  | 2 | MH8919283, MH8919178, MH8919226, MH8919282, MH8919176 |
| Macrophage TREM2<sup>high</sup> | 1 | SA144, SA145, SA222 |
|  | 2 | SA151, SA166, SA227 |
| Macrophage HLA<sup>high</sup>CLEC10A<sup>high</sup> | 1 | SA144, SA145, SA222 |
|  | 2 | SA151, SA166, SA227 |

**Supplementary Table 4.**
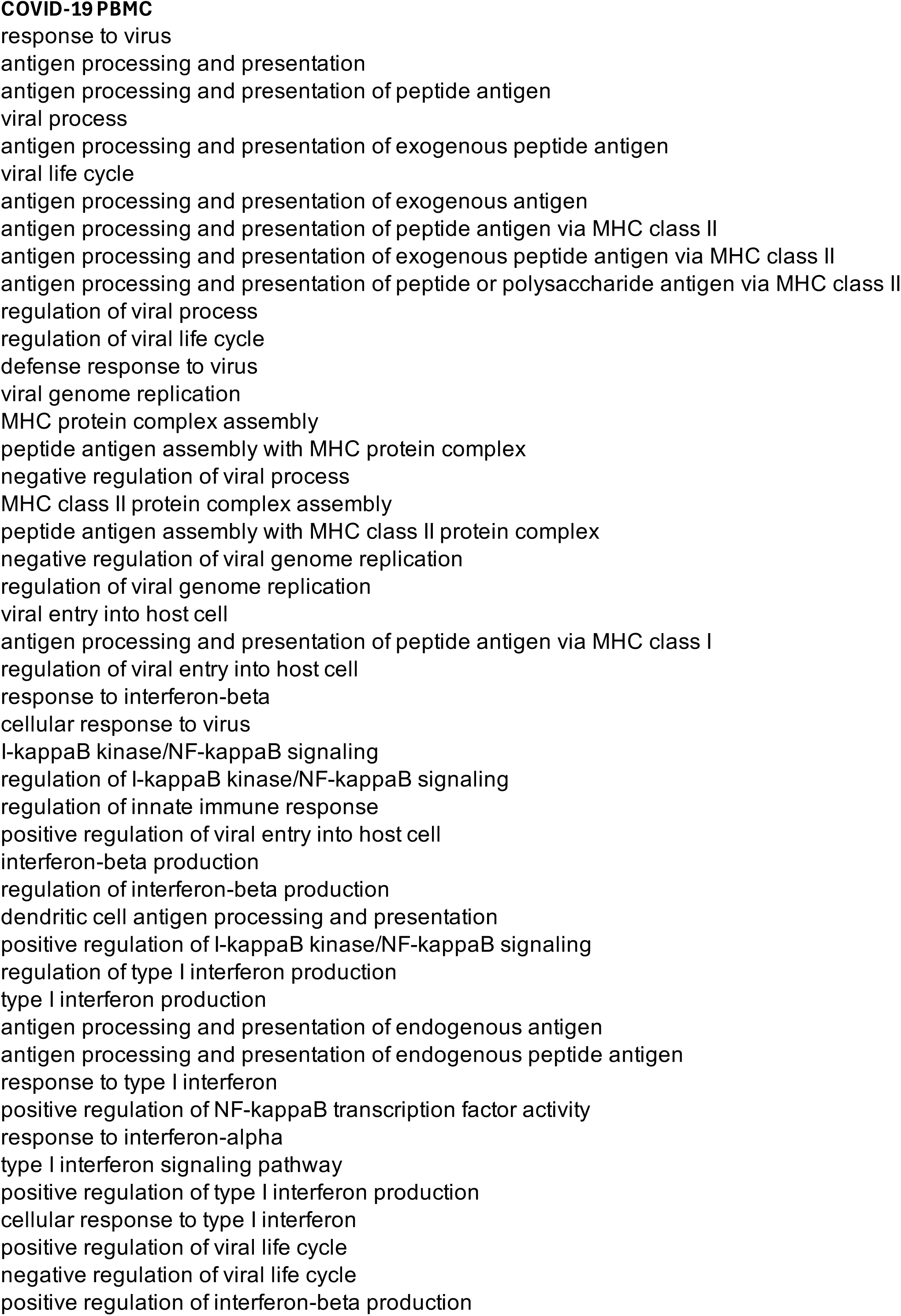
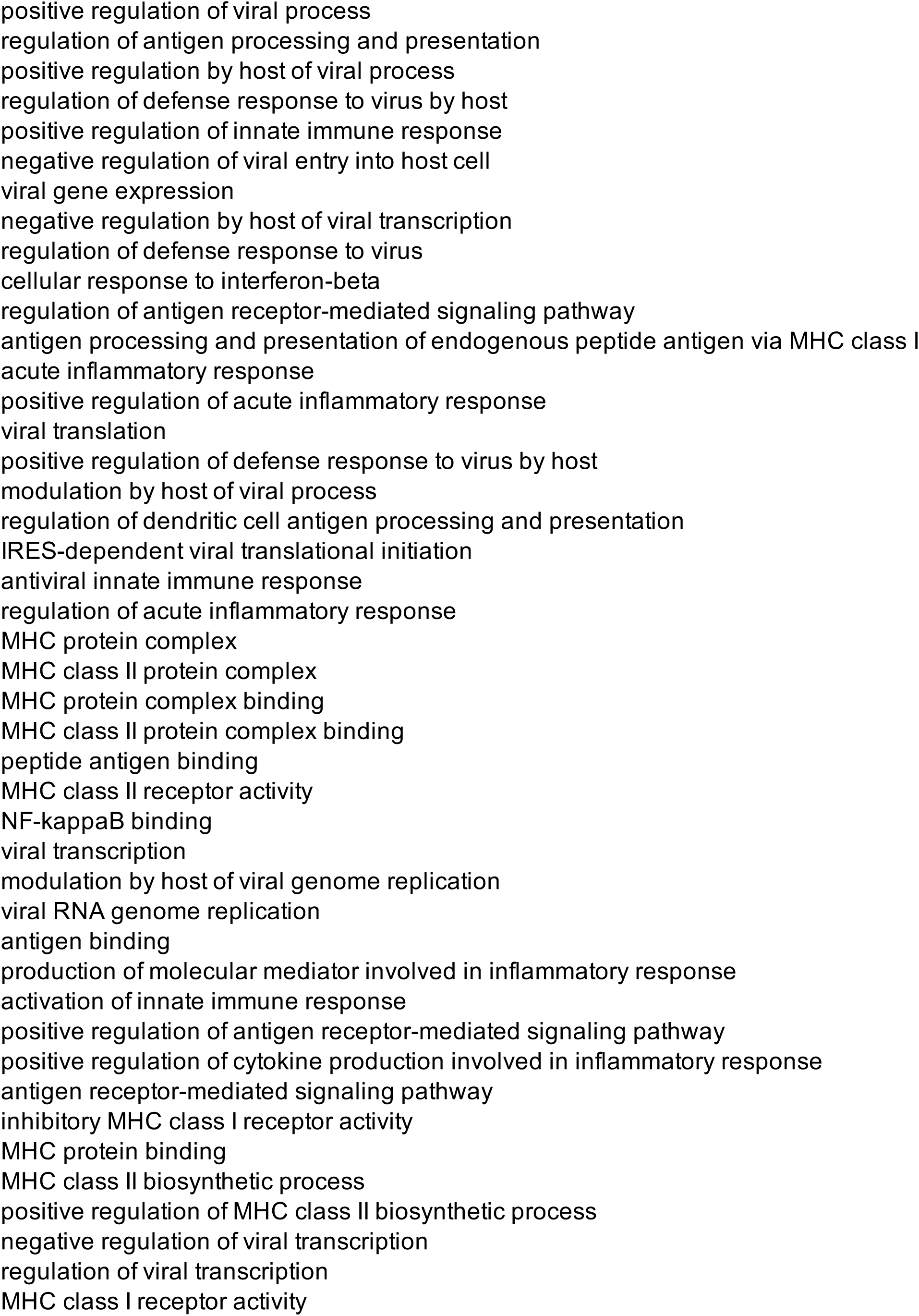
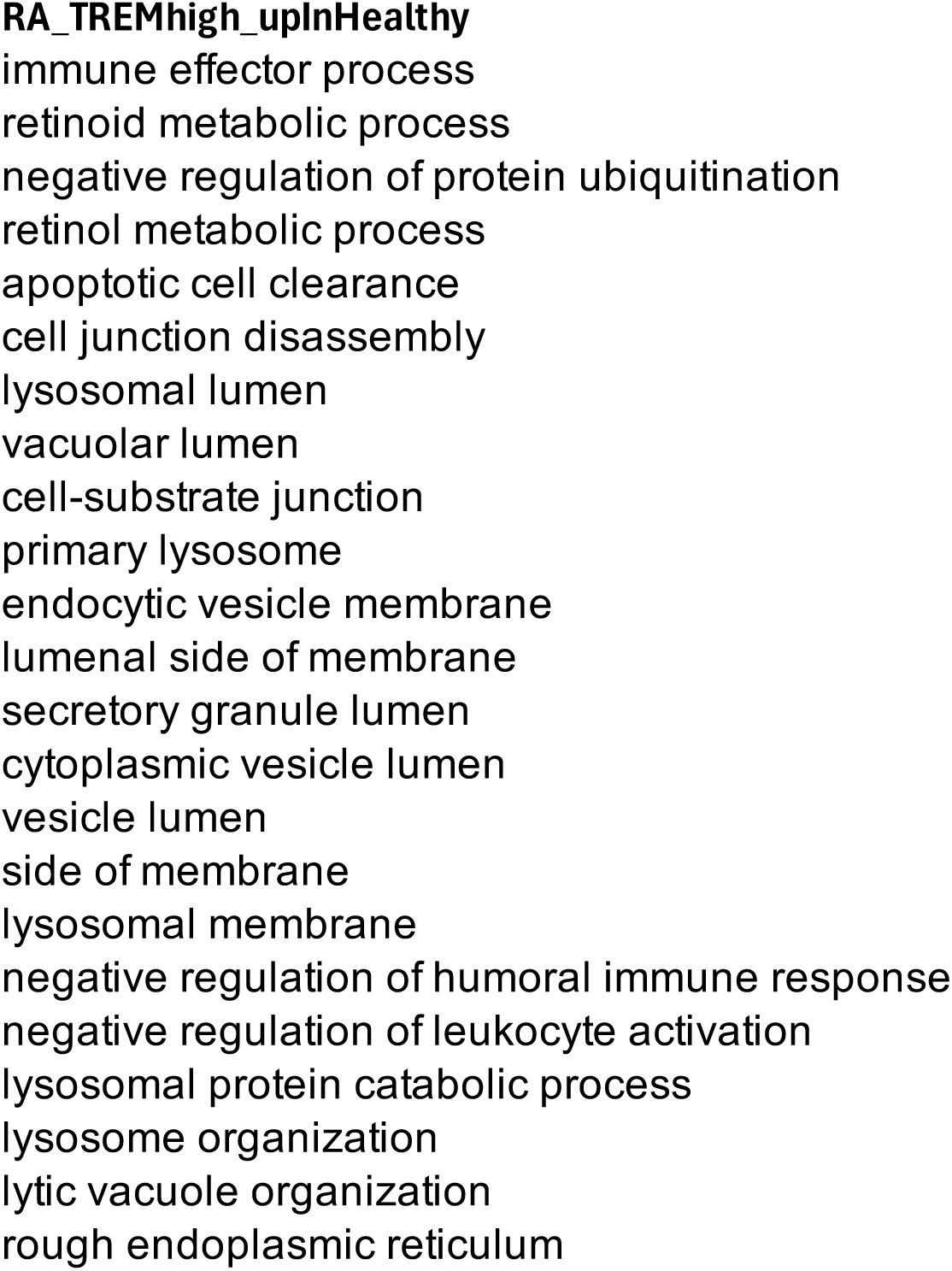
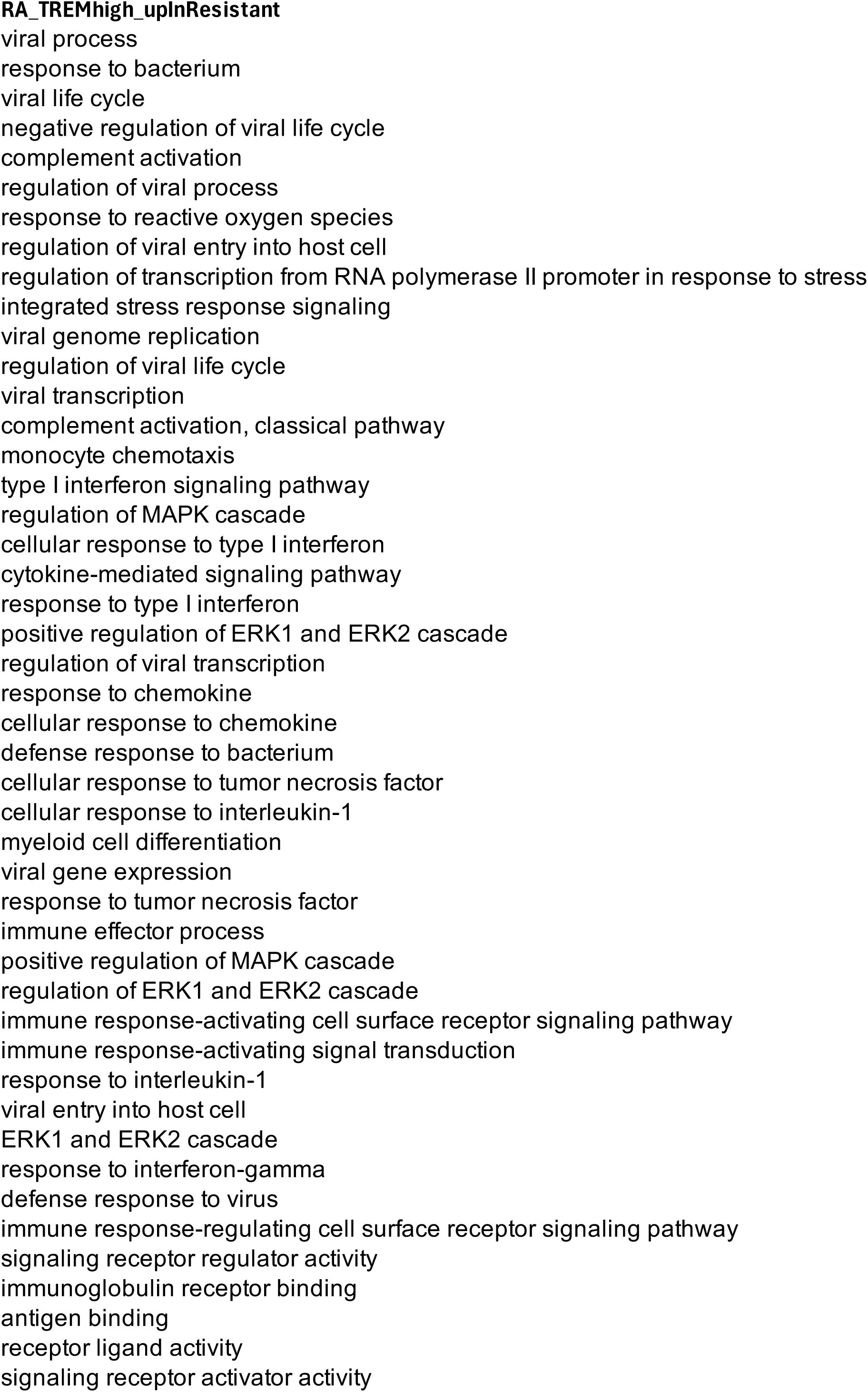
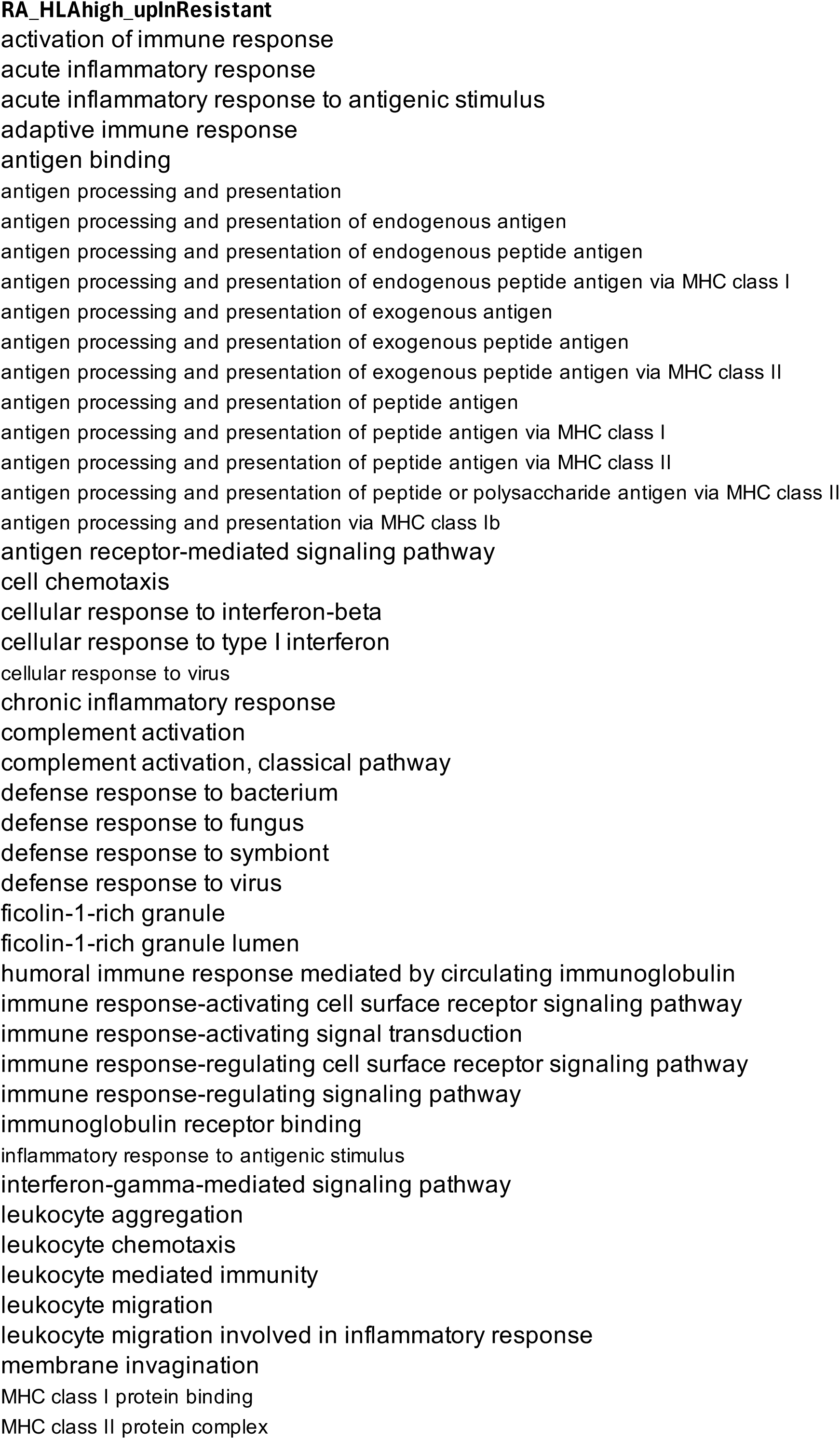
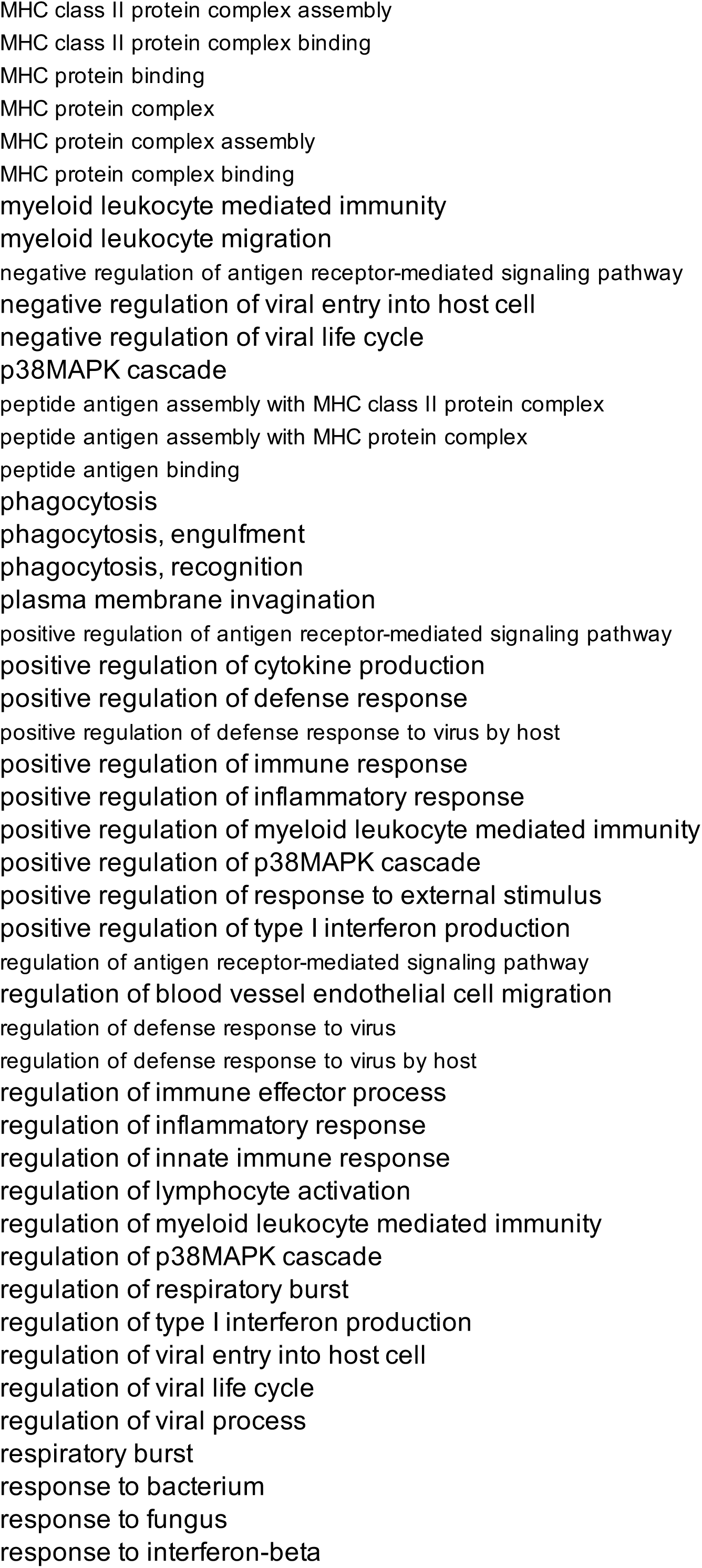
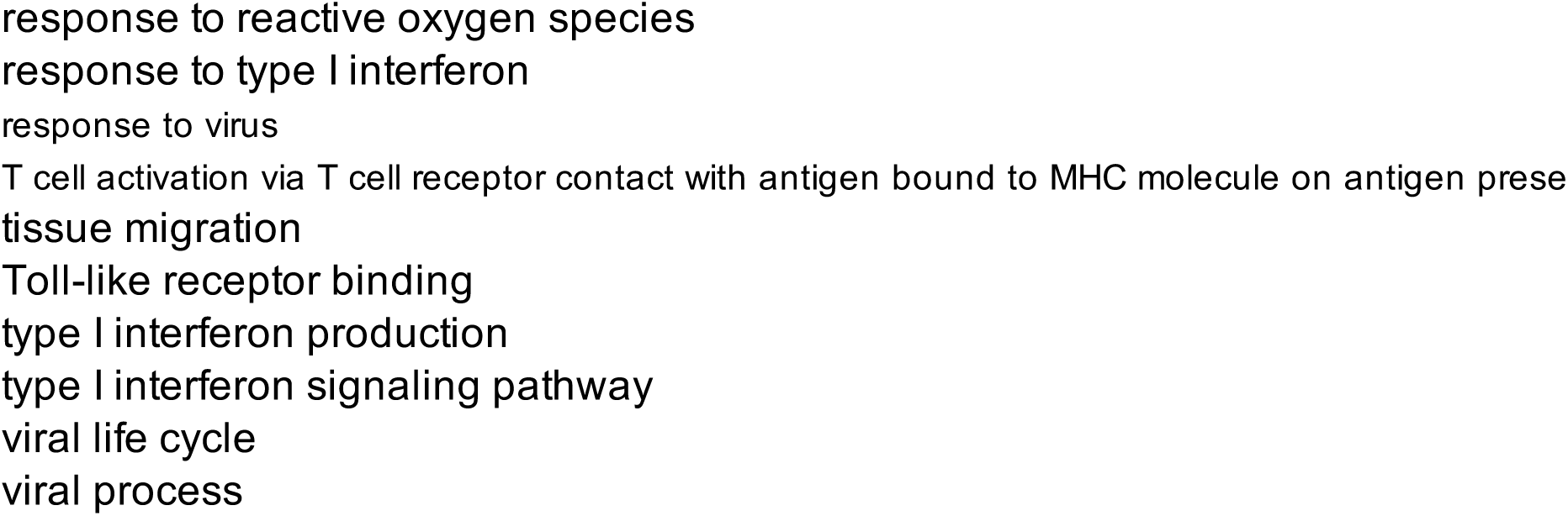
Curated true positive Gene Ontology terms for real dataset evaluation. For each disease and cell type, we curated lists of Gene Ontology (GO) terms expected to be biologically relevant based on established literature. These were used as true positive sets to evaluate method performance in real data. The table includes 91 GO terms related to innate immunity for the COVID-19 dataset, For the rheumatoid arthritis TREM2high cluster, we defined two separate sets: 23 GO terms related to normal tissue homeostasis for genes upregulated in healthy synovial tissue, and 46 GO terms associated with chronic inflammation for genes upregulated in active RA. For the HLAhigh cluster, we focused exclusively on genes upregulated in active RA, selecting 105 GO terms

**Supplementary table 5.** Comparison of CellDEEP strategies for accuracy in Muscat simulation. Read-count aggregation methods (Mean vs. Sum) and cell-selection approaches (K-means vs. Random) are evaluated, result split by DE methods (MAST vs DESeq2). Values represent the median accuracy. 10 cells were aggregated.

| Simulation framework | Aggregation parameter | Cell selection parameter | Accuracy |  |
| --- | --- | --- | --- | --- |
|  |  |  | MAST | DESeq2 |
| Muscat | mean | k-means | 0.95 | 0.96 |
|  |  | Random | 0.95 | 0.96 |
|  | Sum | k-means | 0.96 | **0.99** |
|  |  | Random | 0.97 | **0.99** |

## References

1. Tang, F. et al. mRNA-Seq whole-transcriptome analysis of a single cell. Nat. Methods 6, 377–382 (2009).

2. Jovic, D. et al. Single-cell RNA sequencing technologies and applications: A brief overview. Clin. Transl. Med. 12, e694 (2022).

3. Zimmerman, K. D., Espeland, M. A. & Langefeld, C. D. A practical solution to pseudoreplication bias in single-cell studies. Nat. Commun. 12, 738 (2021).

4. Crowell, H. L. et al. muscat detects subpopulation-specific state transitions from multi-sample multi-condition single-cell transcriptomics data. Nat. Commun. 11, 6077 (2020).

5. Love, M. I., Huber, W. & Anders, S. Moderated estimation of fold change and dispersion for RNA-seq data with DESeq2. Genome Biol. 15, 550 (2014).

6. Robinson, M. D., McCarthy, D. J. & Smyth, G. K. edgeR: a Bioconductor package for differential expression analysis of digital gene expression data. Bioinformatics 26, 139–140 (2010).

7. Ritchie, M. E. et al. limma powers differential expression analyses for RNA-sequencing and microarray studies. Nucleic Acids Res. 43, e47 (2015).

8. Soneson, C. & Robinson, M. D. Bias, robustness and scalability in single-cell differential expression analysis. Nat. Methods 15, 255–261 (2018).

9. Finak, G. et al. MAST: a flexible statistical framework for assessing transcriptional changes and characterizing heterogeneity in single-cell RNA sequencing data. Genome Biol. 16, 278 (2015).

10. Squair, J. W. et al. Confronting false discoveries in single-cell differential expression. Nat. Commun. 12, 5692 (2021).

11. Junttila, S., Smolander, J. & Elo, L. L. Benchmarking methods for detecting differential states between conditions from multi-subject single-cell RNA-seq data. Brief. Bioinform. 23, bbac286 (2022).

12. Nguyen, H. C. T., Baik, B., Yoon, S., Park, T. & Nam, D. Benchmarking integration of single-cell differential expression. Nat. Commun. 14, 1570 (2023).

13. Lee, H. & Han, B. General solution to inflated type I and II errors in multi-subject single-cell differential gene expression analysis. 2023.01.09.523212 Preprint at 10.1101/2023.01.09.523212 (2024).

14. Lee, H. & Han, B. Pseudobulk with proper offsets has the same statistical properties as generalized linear mixed models in single-cell case-control studies. Bioinformatics 40, (2024).

15. Butler, A., Hoffman, P., Smibert, P., Papalexi, E. & Satija, R. Integrating single-cell transcriptomic data across different conditions, technologies, and species. Nat. Biotechnol. 36, 411–420 (2018).

16. Stephenson, E. et al. Single-cell multi-omics analysis of the immune response in COVID-19. Nat. Med. 27, 904–916 (2021).

17. Alivernini, S. et al. Distinct synovial tissue macrophage subsets regulate inflammation and remission in rheumatoid arthritis. Nat. Med. 26, 1295–1306 (2020).

18. Murphy, A. E. & Skene, N. G. A balanced measure shows superior performance of pseudobulk methods in single-cell RNA-sequencing analysis. Nat. Commun. 13, 7851 (2022).

19. Ji, L. et al. The crucial regulatory role of type I interferon in inflammatory diseases. Cell Biosci. 13, 230 (2023).

20. MacDonald, L. et al. Synovial tissue myeloid dendritic cell subsets exhibit distinct tissue-niche localization and function in health and rheumatoid arthritis. Immunity 57, 2843–2862.e12 (2024).

21. Kurowska-Stolarska, M. & Alivernini, S. Synovial tissue macrophages in joint homeostasis, rheumatoid arthritis and disease remission. Nat. Rev. Rheumatol. 18, 384–397 (2022).

22. Song, D. & Li, J. J. PseudotimeDE: inference of differential gene expression along cell pseudotime with well-calibrated p-values from single-cell RNA sequencing data. Genome Biol. 22, 124 (2021).

23. Kim, J. K., Kolodziejczyk, A. A., Ilicic, T., Teichmann, S. A. & Marioni, J. C. Characterizing noise structure in single-cell RNA-seq distinguishes genuine from technical stochastic allelic expression. Nat. Commun. 6, 8687 (2015).

24. Cao, Y., Yang, P. & Yang, J. Y. H. A benchmark study of simulation methods for single-cell RNA sequencing data. Nat. Commun. 12, 6911 (2021).

25. Duo, H. et al. Systematic evaluation with practical guidelines for single-cell and spatially resolved transcriptomics data simulation under multiple scenarios. Genome Biol. 25, 145 (2024).

